# CRUP: A comprehensive framework to predict condition-specific regulatory units

**DOI:** 10.1101/501601

**Authors:** Anna Ramisch, Verena Heinrich, Laura V. Glaser, Alisa Fuchs, Xinyi Yang, Philipp Benner, Robert Schöpflin, Na Li, Sarah Kinkley, Anja Hillmann, John Longinotto, Steffen Heyne, Beate Czepukojc, Sonja M. Kessler, Alexandra K. Kiemer, Cristina Cadenas, Laura Arrigoni, Nina Gasparoni, Thomas Manke, Thomas Pap, Andrew Pospisilik, Jan Hengstler, Jörn Walter, Sebastiaan H. Meijsing, Ho-Ryun Chung, Martin Vingron

## Abstract

We present the software **CRUP** (**C**ondition-specific **R**egulatory **U**nits **P**rediction) to infer from epigenetic marks a list of regulatory units consisting of dynamically changing enhancers with their target genes. The workflow consists of a novel pre-trained enhancer predictor that can be reliably applied across cell lines and species, solely based on histone modification ChIP-seq data. Enhancers are subsequently assigned to different conditions and correlated with gene expression to derive regulatory units. We thoroughly test and then apply **CRUP** to a rheumatoid arthritis model, identifying enhancer-gene pairs comprising known disease genes as well as new candidate genes.

**Availability:** https://github.com/VerenaHeinrich/CRUP

## 1 Background

Gene expression is to a large degree regulated by distal genomic elements referred to as enhancers (Shlyueva *et al.*, 2014), which recruit a combination of different factors to activate transcription from a targeted core promoter. The activity state of enhancers may change dynamically across conditions, e.g. across varying time-points, cell lines, species or disease states. Thus, their activity patterns are central in the context of phenotypic diversity (Wray, 2007; Wittkopp and Kalay, 2012) and altered activity can be the source of pathogenic disruptions of gene-enhancer interactions and subsequent mis-regulation (Maurano *et al.*, 2012). Although the functional importance of enhancers was first observed almost 40 years ago (Banerji *et al.*, 1981), the underlying mechanisms by which enhancers regulate gene expression are still not fully understood. To date, there is neither a complete knowledge of enhancer elements nor their regulatory interplay with targeted genes. However, by analyzing epigenetic profiles of experimentally determined enhancers or binding sites of co-activators like p300 (Visel *et al.*, 2009), condition-specific changes of enhancers were found to be reflected in the epigenetic landscape (Heintzman *et al.*, 2009). Further, to get a glimpse of the underlying causative regulatory mechanism, dynamic enhancer elements need to be further associated with promoter activity across the same conditions. This is particularly important as the majority of gene-enhancer pairs that change dynamically under different conditions have not been discovered, yet (Corradin *et al.*, 2014).

Most experimental procedures which are used to validate regulatory activity of enhancers and their targets are cost and time consuming. Consequently, computational methods that identify enhancer elements based on epigenetic profiles became an indispensable alternative over the last years (Ernst and Kellis, 2012; Mammana and Chung, 2015; He *et al.*, 2017; Rajagopal *et al.*, 2013). However, just a small fraction of enhancers has been functionally characterized in different cell types or tissues (Corradin *et al.*, 2014). Consequently, approaches that rely on a pre-defined gold-standard set of enhancers are often prone to be biased for the cell type or tissue that was used for training. Although strategies that address this shortcoming were recently introduced (He *et al.*, 2017), it remains difficult to develop a method that is able to generalize across different conditions, especially as there are usually just a few common enhancer features available for all data sets. This becomes tremendously important if the overall aim is to identify novel enhancer regions that dynamically change across different conditions. Apart from that, most of the available computational methods are not automatically providing a way to compare many samples across different conditions, and thus the assignment of differential regions has to be done separately in a post-processing step.

Further, the allocation of putative target gene promoters remains challenging, especially as the distance between enhancers and targeted promoters can be very heterogeneous (de Laat and Duboule, 2013). Several methods have already beed introduced that are based on the correlation of epigenomic signals between enhancers and promoters (Corradin *et al.*, 2014; Ernst *et al.*, 2011). One major restriction of these approaches is the missing information about the search space for each enhancer-promoter pair as the linear distance can range from 1kb to several megabases (de Laat and Duboule, 2013). However, this drawback can be compensated by recently introduced methods to determine the direct physical contact between any genomic regions (Dekker *et al.*, 2002; Rao *et al.*, 2014) which also reveal regulatory separated parts of the genome.

In this work, we address all of the above-mentioned issues and present the three-step framework **CRUP** (**C**ondition-specific **R**egulatory **U**nits **P**rediction), that combines the prediction of active enhancer elements (**CRUP-EP**) with a condition-specific assignment (**CRUP-ED**) and the allocation of simultaneously changing gene-enhancer pairs (**CRUP-ET**).

An enhancer classification method that aims to find dynamically changing activity pattern across different conditions needs to be applicable to experiments for which no training data exist. In fact, only a few of the available enhancer prediction tools provide a pre-trained classifier which can be applied to new experimental data. To overcome this bottleneck, we developed the random forest-based enhancer classifier **CRUP-EP** (**E**nhancer **P**rediction) that can be applied across different cell types and species without the need of being re-trained. Here, we made use of the widely accepted concept that enhancer activity is reflected by a certain local chromatin structure with particular histone modifications (HMs) (Creyghton *et al.*, 2010; Rada-Iglesias *et al.*, 2011), which can be determined by ChIP-seq (Horak and Snyder, 2002). The local structure comprises in essence an accessible region flanked by nucleosomes which carry a H3K27ac modification and where the H3K4me1 signal dominates over H3K4me2 or -me3. Moreover, the proportion of H3K4me1 over H3K4me3 distinguishes an enhancer from a promoter region (Heintzman *et al.*, 2007). Based on these characteristics **CRUP-EP** solely requires the six core HMs on which the NIH Roadmap Epigenomics Mapping Consortium (Bernstein *et al.*, 2010) and the International Human Epigenome Consortium (IHEC, Stunnenberg *et al.* 2016) have converged, guaranteeing a broad applicability. We train and validate **CRUP-EP** on mouse embryonic stem cells (mESCs) based on curated FANTOM5 validated enhancer regions (de Hoon *et al.*, 2010) such that our training and test sets are chosen independently from the HM feature set that is used as input for our enhancer classifier. To validate the open chromatin property, we use the distance to the nearest accessible region as an additional quality measure for our predicted enhancer regions by integrating an independent ATAC-seq experiment (Buenrostro *et al.*, 2015).

Recently, super enhancers (SEs) were introduced as an important subset of large (> 3 kb) enhancer regions, which are especially crucial for the regulation of expression of cell identity genes (Whyte *et al.*, 2013). Hence, to evaluate our predictions, we further cluster single enhancers in mESCs and show that almost all of these are overlapping with a list of SEs recently published by Novo *et al.* (2018).

To demonstrate the transferability of **CRUP-EP** we integrated five different experimental data sets, comprising various cell types and species, that were obtained in the context of the German Epigenome Project (DEEP, 2017). We trained and applied our classification approach on mESC as well as on the DEEP-related data sets and validated the performance across the different samples. We further compare our results to two other widely used enhancer prediction methods, namely ChromHMM (Ernst and Kellis, 2012) and REPTILE (He *et al.*, 2017). Hidden Markov Models (HMMs), integrated e.g. in ChromHMM, are well suited to discover unknown combinations of chromatin signatures from HM ChIP-seq experiments and especially ChromHMM is widely used when it comes to enhancer prediction. HMM-based approaches usually do not include prior knowledge into the predictions but the interpretation of the segmentation and the choice of the number of chromatin states has to be done manually. On the other hand, REPTILE implements a random forest-based algorithm to assign enhancer probabilities to each region in the genome and is trained on distal binding sites of the histone modifying protein p300. In this work we refrain from further method comparisons since REPTILE just recently demonstrated to be superior to several state-of-the-art enhancer prediction tools, e.g. RFECS (Rajagopal *et al.*, 2013). We show that **CRUP-EP** outperforms ChromHMM and is comparable to REPTILE when applying within a single cell type. In terms of transferability across different cell types and species, we can show that our classification approach outperforms REPTILE.

A prominent application of enhancer prediction methods is be the comparison of dynamic conditions, like varying time-points, cell lines or disease states. To address this, we complement **CRUP-EP** by **CRUP-ED** (**E**nhancer **D**ynamics) which assigns predicted enhancer regions to distinguishable conditions while accounting for a flexible number of replicates. Based on the enhancer probabilities obtained by **CRUP-EP**, the second phase of **CRUP** is computing pair-wise empirical p-values based on a permutation test which are further used to cluster significantly different enhancer regions.

We apply **CRUP-ED** to a data set of pluripotent and retinoic acid (RA) induced mESCs yielding two clusters of condition-specific enhancer regions. We can show that the assignment of clusters across the two conditions is in good agreement with HM ChIP-seq as well as with ATAC-seq data. Enhancer elements regulate gene expression through the binding of sequence-specific transcription factors (TFs) to cognate motifs (Grossman *et al.*, 2017). Therefore, we further evaluate our dynamic enhancer regions by investigating the over-representation of TF motifs within each enhancer cluster. We are able to identify several motifs that are associated with RA receptors as well as with signaling pathways that regulate the pluripotency of stem cells.

A traditional approach to infer a gene promoter that is targeted by an enhancer is to apply a distance-based strategy whereas either the nearest promoter is chosen or some statistic is applied within a fixed search space. However, it has been recently shown that enhancer activation can not only skip several non-target genes, independently of their orientation, but enhancer elements can also be placed in the gene body of another independent gene (Pennacchio *et al.*, 2013).

To understand the underlying regulatory mechanism of dynamic enhancer regions, they need to be linked to promoter activity across the same conditions. As enhancer dynamics strongly correlate with changing gene expression pattern (Heintzman *et al.*, 2009), we make use of this property and added a third layer to our framework, **CRUP-ET** (**E**nhancer **T**argets), to match condition-specific enhancers to RNA-seq experiments (Wang *et al.*, 2009).

Recently, conformation capture methods such as Hi-C (Rao *et al.*, 2014; Bonev *et al.*, 2017) or CaptureC-seq (Andrey *et al.*, 2017) have focused on the three-dimensional non linear structure of the genome. Chromatin folding brings distal regulatory elements, such as enhancers, into close physical proximity of their target gene promoters (Tolhuis *et al.*, 2002). Although, until now, single enhancer-promoter contacts have been difficult to observe in Hi-C experiments, they have been shown to interact primarily within the same topological associated domain (TAD) (Bonev *et al.*, 2017; Yin *et al.*, 2012). In this work we make use of this knowledge and designed **CRUP-ET** in such a way that it restricts the search space to prioritize promoter/gene-enhancer interactions within the same TAD.

We identify differential enhancer regions across eight developmental states in mouse embryo mid-brain and link each region to putative target genes. Using a small set of genes that are found to be active in the same tissue we show that our inferred regulatory units (high-confidence gene-enhancer pairs) coincide well with corresponding CaptureC-seq data and recapitulate physical interaction information. We further evaluate our approach using a data set comprising three states of mouse neural differentiation and demonstrate a very good agreement of our inferred dynamic regulatory units using ultra-deep Hi-C data.

Finally, we identify trait-associated regulatory elements in a mouse model of rheumatoid arthritis, an autoimmune inflammatory complex disease, and discuss our main findings on a single enhancer region that we can correlate to several genes of the *CCR*-gene cluster, which is part of the Chemokine signaling pathway. Additionally, we support our findings with motif enrichment analysis as well as with a pathway analysis. With this, we demonstrate how our presented framework **CRUP** can be used to identify candidate enhancer regions together with their putative target genes that dynamically change between different conditions.

## 2 Results

### 2.1 Short summary of CRUP

In this work, we describe the three-step framework **CRUP** to predict active enhancer regions, assign them to specific conditions and finally correlate each dynamically changing enhancer to putative target genes. Each single step is implemented in R and incorporated into a workflow (Figure 1). The first module of our framework, **CRUP-EP** (**E**nhancer **P**rediction, Section 5.8), is an enhancer classifier with feature sets based on six core HMs, namely H3K4me1, H3K4me3, H3K27ac, H3K36me3, H3K27me3 and H3K9me3 (Figure 1 A). We implemented a combination of two random forests to split the task of distinguishing active regulatory regions from the rest of the genome, as well as differentiating enhancers from active promoters. **CRUP-EP** is designed such that it takes into account the basic genomic structure of an enhancer, which is in essence an open chromatin region flanked by nucleosomes.

**Figure 1:**
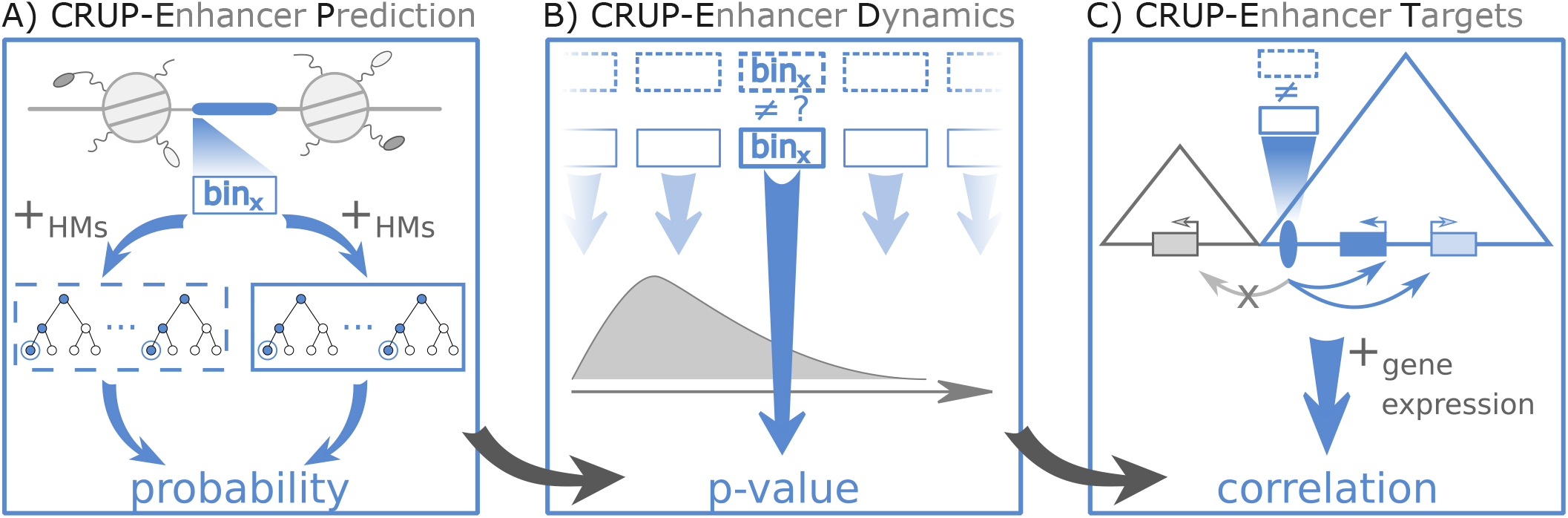
Schematic overview. **CRUP** (**C**ondition-specific **R**egulatory **U**nits **P**rediction) is a three-step framework to predict active enhancers (’*CRUP-EP*’), assign them to dynamic conditions (’*CRUP-ED*’) and create differential regulatory units (’*CRUP-ET* ’). **A)** CRUP-EP accounts for the size of accessible regions (highlighted in blue) which are flanked by nucleosomes. For each region of interest, bin_*x*_, a combination of two binary random forest classifiers, solely based on ChIP-seq HM data, is then used for enhancer prediction. **B)** Based on a permutation test, CRUP-ED computes empirical p-values for each bin_*x*_ across different conditions (dotted and solid rectangles), which are further used to combine and cluster regions. **C)** CRUP-ET inspects each differential enhancer region (blue ellipse) individually within its topologically associated domain (blue triangle). To infer putative target genes, the correlation between probability values and gene expression counts is calculated.

The second phase of the workflow, **CRUP-ED** (**E**nhancer **D**ynamics, Section 5.9), is based on genome-wide enhancer predictions for multiple conditions, e.g. different development states of a cell (Figure 1 B). We find condition-specific enhancers by applying a permutation test directly on the predicted enhancer probabilities obtained by **CRUP-EP**. Based on pairwise empirical p-values, dynamic enhancers can be combined and clustered into differential regions.

In a last step, **CRUP-ET** (**E**nhancer **T**argets, Section 5.10), each dynamically changing enhancer region obtained by **CRUP-ED** is linked to target genes (Figure 1 C). To this end, the correlation between enhancer probabilities and gene expression values across the same conditions is computed for all putative gene-enhancer pairs that are located within the same TAD.

We trained **CRUP-EP** on input-normalized HM ChIP-seq data and a training set based on FANTOM5 curated enhancers. To evaluate **CRUP-ED** and **CRUP-ET** we predicted active enhancer regions based on a classifier trained on mouse embryonic stem cells (mESC).

### 2.2 Accuracy and spatial resolution of enhancer predictions in mESC

We trained our random forest based enhancer classifier **CRUP-EP** on input-normalized HM ChIP-seq data from a single mESC sample, in this work further labeled as mESC^+^ (see Section 5.8). The predictions were validated on ten test sets, primarily focusing on the area under the precision-recall (AUC-PR) curve. Overall, our classification method yields very good and stable results across all test sets with an AUC-PR ranging from 0.93 to 0.96 and an AUC-ROC ∈ [0.98, 0.99] (Figure 2 A, Figure S2). This also holds true when comparing to two other widely used enhancer classification methods, ChromHMM and REPTILE, which will be further discussed in Section 2.4.

**Figure 2:**
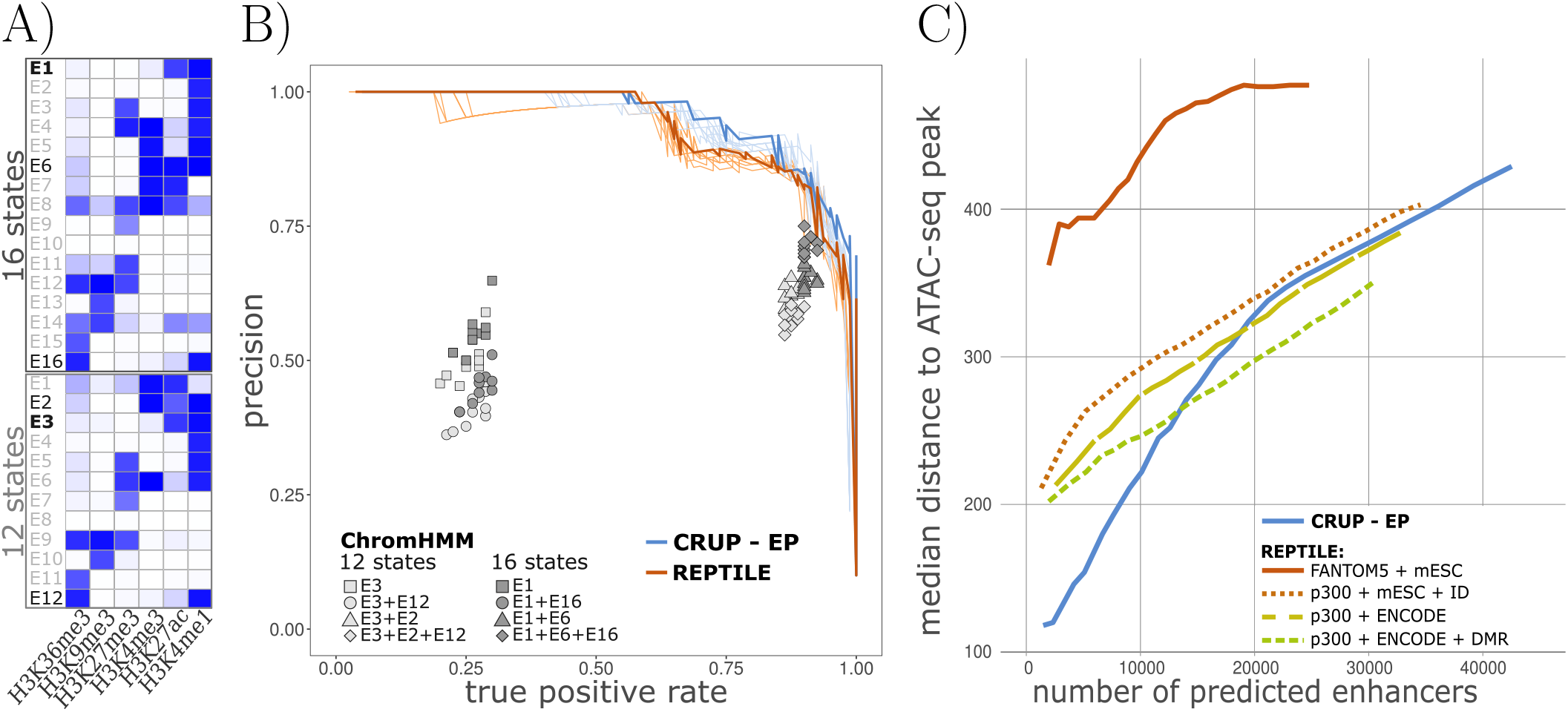
Performance of enhancer classifiers in murine ESC. **A)** ChromHMM emission probabilities for mESC using 12 and 16 chromatin states, ranging from 0 (white) to 1 (dark blue). The state that resembles enhancers best is marked as bold. **B)** Precision recall curves for CRUP-EP (light blue lines) and REPTILE (light orange lines) trained on an mESC sample (mESC^+^) and tested on ten randomly sampled independent test sets. The curves for the best performances are highlighted in darker colors. Additionally, the performance results of different ChromHMM segmentations for the same ten test sets are depicted (gray shapes). **C)** The median distance of predicted mESC enhancers to the closest ATAC-seq peak for CRUP-EP (blue) and different settings of REPTILE. The evaluated sets of predicted enhancers are based on decreasing probability cutoffs ∈ [1, 0.5]. REPTILE was trained on our FANTOM5-based training set and mESC^+^ (orange), on a p300-based training set, mESC^+^ and intensity deviation (ID) features (dotted light orange), on a p300-based training set using mESC ENCODE data (dashed yellow) and on a p300-based training set using mESC ENCODE data and differentially methylated regions (DMRs, dashed green).

Based on the genome-wide predictions we called 42, 530 enhancers at length 1100 bp, using a minimum probability threshold of 0.5 as described in Section 5.8. The distribution of the original HM ChIP-seq read counts over the called enhancers shows a partition of these regions into a large and a small cluster (Figure S5). The majority of the predicted enhancers show typical enhancer characteristics, with high enrichment for the histone marks H3K4me1 and H3K27ac, as well as for an independent ATAC-seq experiment. A much smaller cluster rather looks like promoter proximal enhancers with an additional, not centered, enrichment for the promoter mark H3K4me3 and an ATAC-seq profile showing several peaks of enrichment at the predicted enhancer. Nevertheless, we do not see any promoter proximal enhancers when applying a more stringent cutoff (0.98) to the probabilities (see Figure S6). To validate the spatial resolution of our predictions, we computed the distance between each of our predicted enhancers and the closest accessible region measured with ATAC-seq (see Section 5.5). By decreasing the probability cutoff from 1 to 0.5 in a step-wise manner, we get an increasing set of enhancers for which we computed the median distance to the closest ATAC-seq peak (Figure 2 C). The top-ranked enhancers show a very low median distance of 118 bp, and for the top ~ 10, 000 regions (probability threshold of 0.84) we still observe a very good resolution with a median distance of 222 bp.

We further investigated the length and distribution of our enhancer predictions in the genome by dividing the whole set into 9, 170 singletons and 7, 550 enhancer clusters covering multiple closely placed enhancer peaks (see Section 5.8) for which two examples are depicted in Figure S7. We compared each enhancer cluster to a list of 927 super-enhancers (SEs) which was recently published by Novo *et al.* (2018). Over 96% (896) of the SEs overlap with our enhancer clusters and almost all of them overlap with our complete non-clustered list of predicted peaks (924/42, 530). This shows that **CRUP-EP** is well suited to capture enhancer regions of very heterogeneous lengths.

For further validation we used a probability cutoff of 0.5 which is implemented by default in **CRUP-EP** to define enhancer regions.

### 2.3 Enhancer predictions are stable across different cell types and species

We trained our enhancer classification approach **CRUP-EP** for 12 different samples from different cell types and species (summarized in Table S2) in the same fashion as described for mESC^+^ in Section 2.2. We used each of the classifiers to predict active enhancers on the test sets of the remaining 11 samples and calculate the AUC-PR, resulting in a 12 × 12 AUC-PR-matrix which is summarized in Figure 3 A. Within one sample, training and test sets are independent following the logic described in Section 5.8.

**Figure 3:**
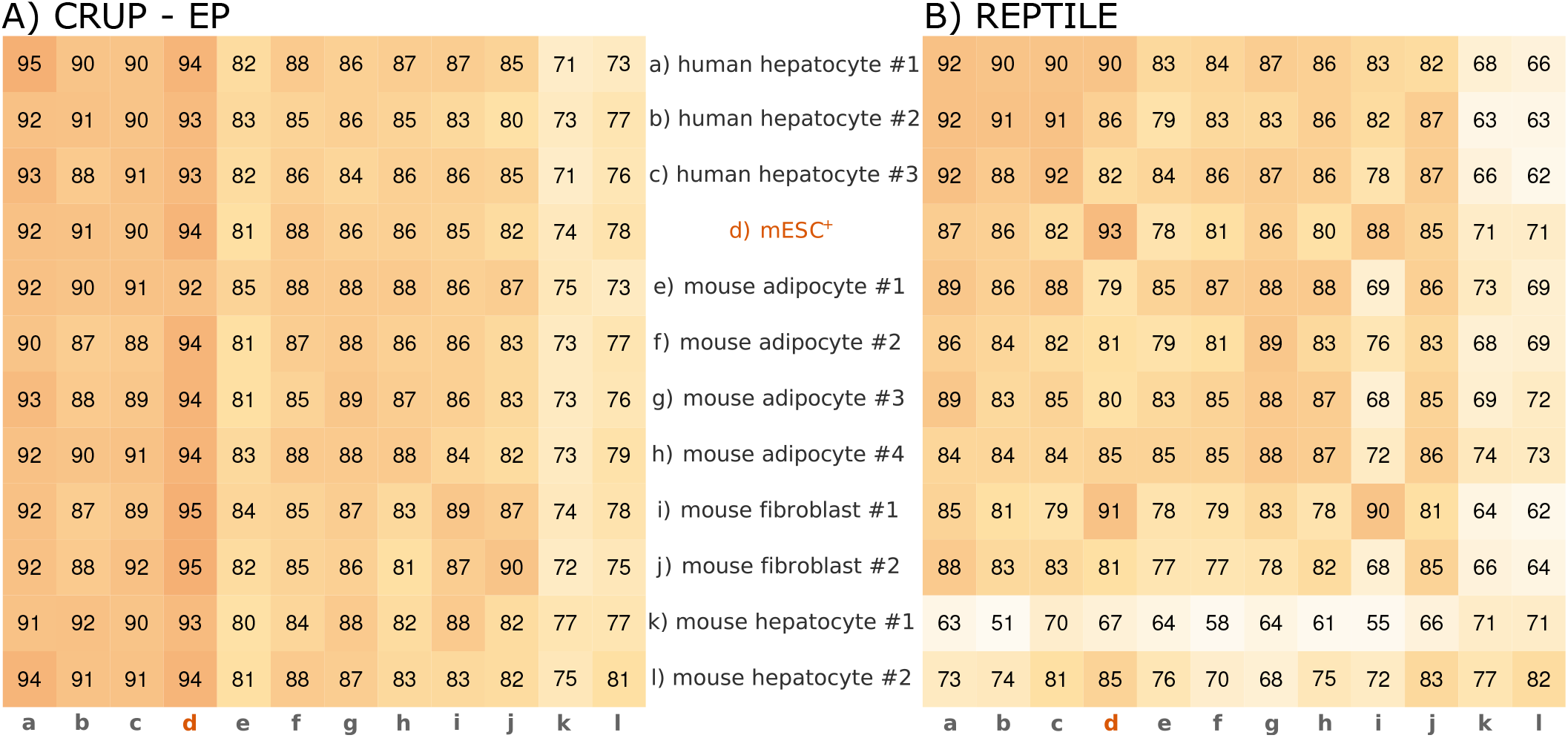
Predictions across different cell types and species. Classifiers were trained on and applied to samples from different cell types (hepatocyte, ESC, adipocyte, fibroblast) and species (mouse and human). The results for CRUP-EP (A) and REPTILE (B) can be summarized in 12 × 12 heatmaps where each entry is shaded according to the computed AUC-PR (in percent). The origin of the training data a)-l) can be found in the rows and the origin of the test sets a-l in the columns. The diagonal shows the performance results on an independent test set within one sample. For instance, using CRUP in mESC^+^ (training and test set highlighted in red) led to an AUC-PR= 0.94.

All classifiers perform well regardless of the test set they are applied to (AUC-PR ∈ [0.71, 0.95]). Interestingly, the performances correlate much more with the test set than with the training set origin, as can be observed in a vertical trend of the AUC-PR values in Figure 3 A. For instance, the lowest AUC-PR value with a minimum of 0.71 is achieved when using one of the mouse hepatocyte samples as a test set. On the other hand, when training the classifier on any mouse hepatocytes sample and testing on a high quality sample, such as mESC^+^, the performance is very good (AUCPR ∈ [0.93, 0.94]). Also, training and prediction within one sample (diagonal entries) rarely results in the best prediction performance for the corresponding classifier. Overall, the best performances across all cell types and species could be achieved when testing on the mESC^+^ sample (AUC-PR ∈ [0.92, 0.95]) and this will be used as the pre-trained classifier provided in **CRUP-EP**.

### 2.4 Comparison to other enhancer prediction methods

Here we compare our enhancer classification approach **CRUP-EP** to two other widely used methods, namely ChromHMM and REPTILE. First, we applied both methods to the undifferentiated mESC^+^ sample and compared the results to the performance of **CRUP-EP** as described in Section 2.2. A more detailed description of the implementation of both methods can be found in Section 5.11.

We created three genome-wide segmentations utilizing ChromHMM with different numbers of chromatin states, *K* ∈ {8, 12, 16}, and defined enhancer states based on the parameters of the emission distribution (Figure 2 A and Figure S8). We validated the segmentation results based on *K* = 12 and *K* = 16 and different combinations of possible enhancer states on ten test sets leading to strongly varying true positive rates (TPRs) ∈ [0.2, 0.925] and precision values ∈ [0.36, 0.775] (Figure 2 B). Compared to the **CRUP-EP** results, the performance of ChromHMM is less robust and less stable. Interestingly, the results cluster into two distinct groups, depending on whether the enhancer definition is only based on high emission probabilities for H3K4me1 and H3K27ac (*E*3 for *K* = 12, *E*1 for *K* = 16) or additionally on the promoter mark H3K4me3 (*E*2 for *K* = 12, *E*6 for *K* = 16). These findings show that there is no state in the ChromHMM segmentations that uniquely describes enhancers or promoters and illustrate the difficulties in separating these two regulatory elements. When we trained REPTILE on our FANTOM5-based training set and six core HMs, we observe a similar but slightly worse test set performance in mESC^+^ (AUC-PR ∈ [0.92, 0.94]) compared to our classifier (AUC-PR ∈ [0.93, 0.96]), Figure 2 A, Figure S2). The differences in performance become more prominent when comparing different training and test set combinations across cell types and species as described in Section 2.3. For most of the combinations of different training and test sets, our classifier (AUC-PR ∈ [0.71, 0.95]) outperforms REPTILE (AUC-PR ∈ [0.51, 0.93]) as depicted in Figure 3. In addition, we used the REPTILE classifier close to its original setting, i.e., trained on the mESC HM features and the p300-based enhancers as described in He *et al.* (2017), to make predictions across the 12 samples. This lead to slightly worse results on the FANTOM5-based test set than when trained on our data (Figure S9).

Next, we measured the median distance to the closest ATAC-seq peak (spatial resolution) of the REPTILE predictions in mESC^+^ following the same procedure as in Section 2.2. To this end, we applied REPTILE in four different training set and feature set combinations. For the training set, we used either FANTOM5-based or p300-based enhancers, and for the feature sets we used our six core HMs, the HM data and the differentially methylated regions (DMRs) from He *et al.* (2017) as well as additional intensity deviation features (see Section 5.11 for more details). Only when including DMRs, REPTILE achieves better results for more than the top 12, 500 predicted enhancers compared to **CRUP-EP** (Figure 2 C). Without DMRs as additional features, **CRUP-EP** performs similar to REPTILE when using p300-based enhancers as a training set. However, when applying the same feature and training set combination, **CRUP-EP** outperforms REPTILE in terms of spatial resolution.

### 2.5 Enhancer probabilities enables identification of clusters of differential enhancers

We applied our approach to identify differential enhancers, **CRUP-ED**, between murine pluripotent (mESC^+^) and differentiated retinoic acid (RA) induced stem cells (mESC^−^) as further described in Section 5.9. To this end, enhancer prediction was performed on both samples using **CRUP-EP** which was trained on mESC^+^ as described in Section 5.8. Dynamically changing enhancer regions that are either active in mESC^+^ (cluster 1) or in the RA-induced mESC^−^ sample (cluster 2) were identified and further summarized as explained in Section 5.9. From the predicted condition-specific enhancers, a total of 58 are only active in mESC^+^ (cluster 1) and 54 regions are predicted to be active solely in mESC^−^ (cluster 2). The differential assignment of predicted enhancers can be further corroborated by ChIP-seq read count distributions (Figure 4 B, also shown for a single differential region in Figure 4 C). The signal for the enhancer marks H3K27ac and H3K4me1 is higher in mESC^−^ (orange) compared to mESC^+^ (gray) for the displayed regions in cluster 2. The same trend can also be observed when investigating chromatin accessibility for the two data sets which becomes detectable via additional ATAC-seq experiments (right panel Figure 4 B, bottom panel 4 C). To further evaluate the two differentially active enhancer clusters, we performed a motif enrichment analysis for both groups as described in more detail in Section 5.12, taking the union of all differential enhancers as the basis for the estimation of the background model. The complete list of differentially enriched motifs is depicted in Figure S13.

**Figure 4:**
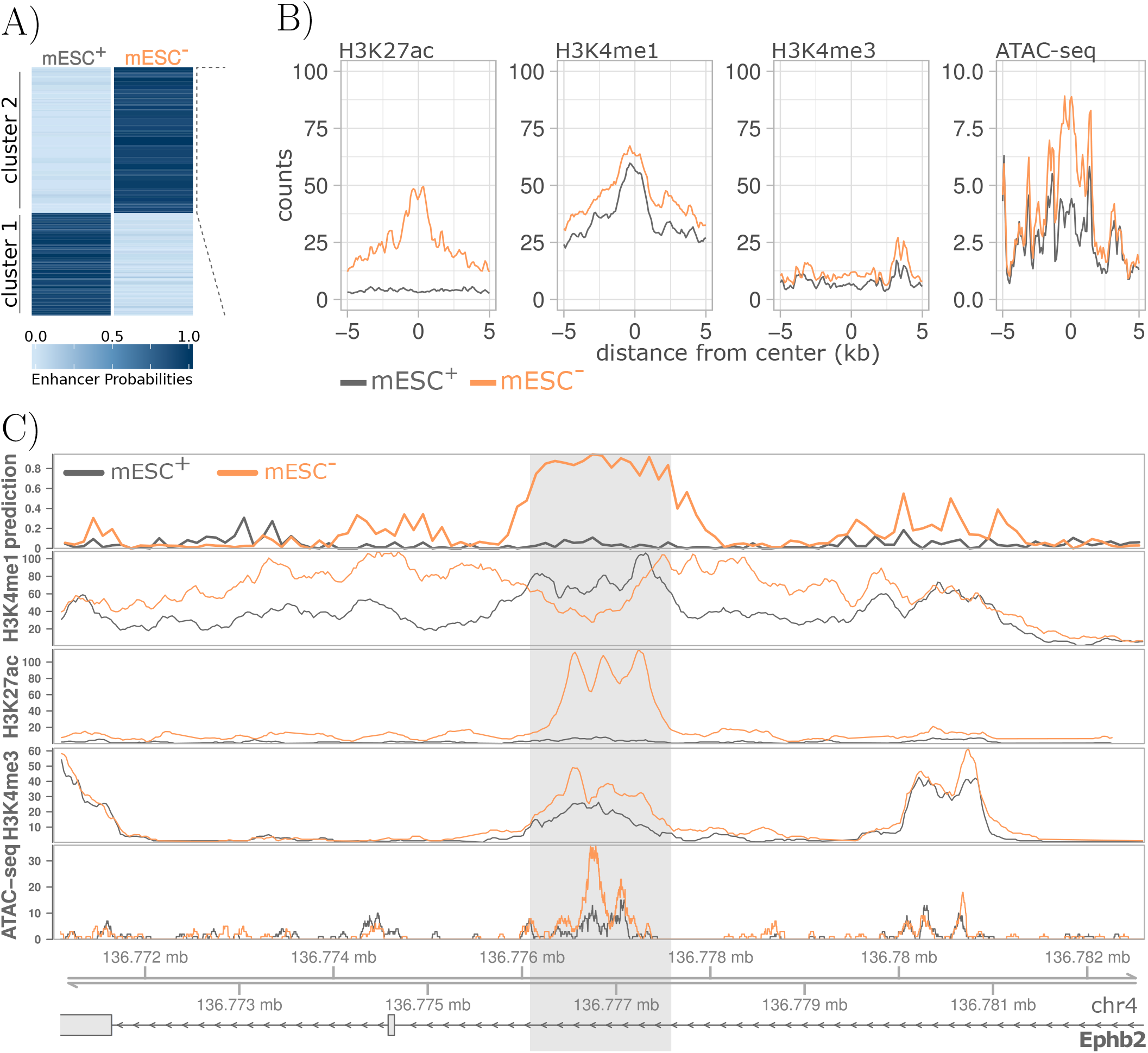
Differential enhancers in murine stem cell differentiation. **A)** Differential enhancer regions of undifferentiated (mESC^+^) and differentiated (mESC^−^) cells, colored by their respective enhancer probabilities. All regions can be divided into two clusters according to their differential activity pattern. **B)** Density distributions of HM ChIP-seq and ATAC-seq read counts for all differential enhancer regions for mESC^+^ (gray) and mESC^−^ (orange) in cluster 2. **C)** The enhancer region (*chr4:*136, 776, 101 − 136, 777, 600, highlighted in light gray) was predicted to be active in mESC^−^ (orange) but not in mESC^+^ (gray). This trend is also apparent by visual inspection of the enhancer probabilities displayed in the top panel (’*prediction*’). The region is located within the intronic region of the gene Ephb2 (blue) which was identified as regulator of stem and progenitor cell proliferation.

With the functional annotation tool DAVID (Huang and Lempicki, 2009; Huang *et al.*, 2009) we could identify several transcription factors (TFs) that show a higher binding site enrichment in cluster 1 and are part of signaling pathways regulating pluripotency of stem cells (OCT4, HNF1A). In the same way, TFs that are more enriched in the second (RA-specific) cluster were found to be linked to the functional categories *’differentiation’* and/or *’developmental protein’* (GLIS2, ASCL1, INSM1, Myod1, Myog, NHLH1, NR2C2, PAX5). Furthermore, we found retinoic acid receptors (heterodimers) in our list of differential transcription factor binding sites (TFBSs) for cluster 2 (RARA::RXRG, RARA::RXRA, Pparg::RXRA, Gudas and Wagner (2011); Cunningham and Duester (2015); Lin *et al.* (2011)).

Binding sites for two differentiation associated TFs with a very similar motif (Myog and Myod1; pvalue ≤ 0.05, see Section 5.12) can be found in a differentially active enhancer region located in an intron of the gene *Ephb2* (Figure 4C). Eph receptors constitute the largest subgroup of tyrosine kinase receptors which were already identified as regulators of stem and progenitor cell proliferation in mice (Chumley *et al.*, 2007). Moreover, *Ephb2* has also been reported to be regulated by retinoic acid (RA) signaling in the chick retina (Sen *et al.*, 2005).

### 2.6 Dynamic enhancers can be linked to putative target genes by a greedy matching approach

By including RNA-seq experiments (see Section 5.3), we utilize **CRUP-ET** to link dynamically changing enhancers to putative target genes (Section 5.10) using a *’greedy matching’* approach. To do so, we calculate Pearson’s correlation coefficients between enhancer probabilities of a differential enhancer region across all samples and normalized expression counts of genes that are located within the same TAD. We further describe dynamically changing gene-enhancer pairs with a correlation coefficient ≥ 0.9 as *’regulatory units’*.

We applied **CRUP-EP** and **CRUP-ED** to predict enhancers and assign them to different conditions in a time series experiment performed in mouse embryo midbrain, spanning eight time points in total (Gorkin *et al.*, 2017). This results in 3, 815 differentially active enhancers that could be grouped and summarized into 379 different clusters using activity pattern as described in Section 5.9 (Figure 5 A).

**Figure 5:**
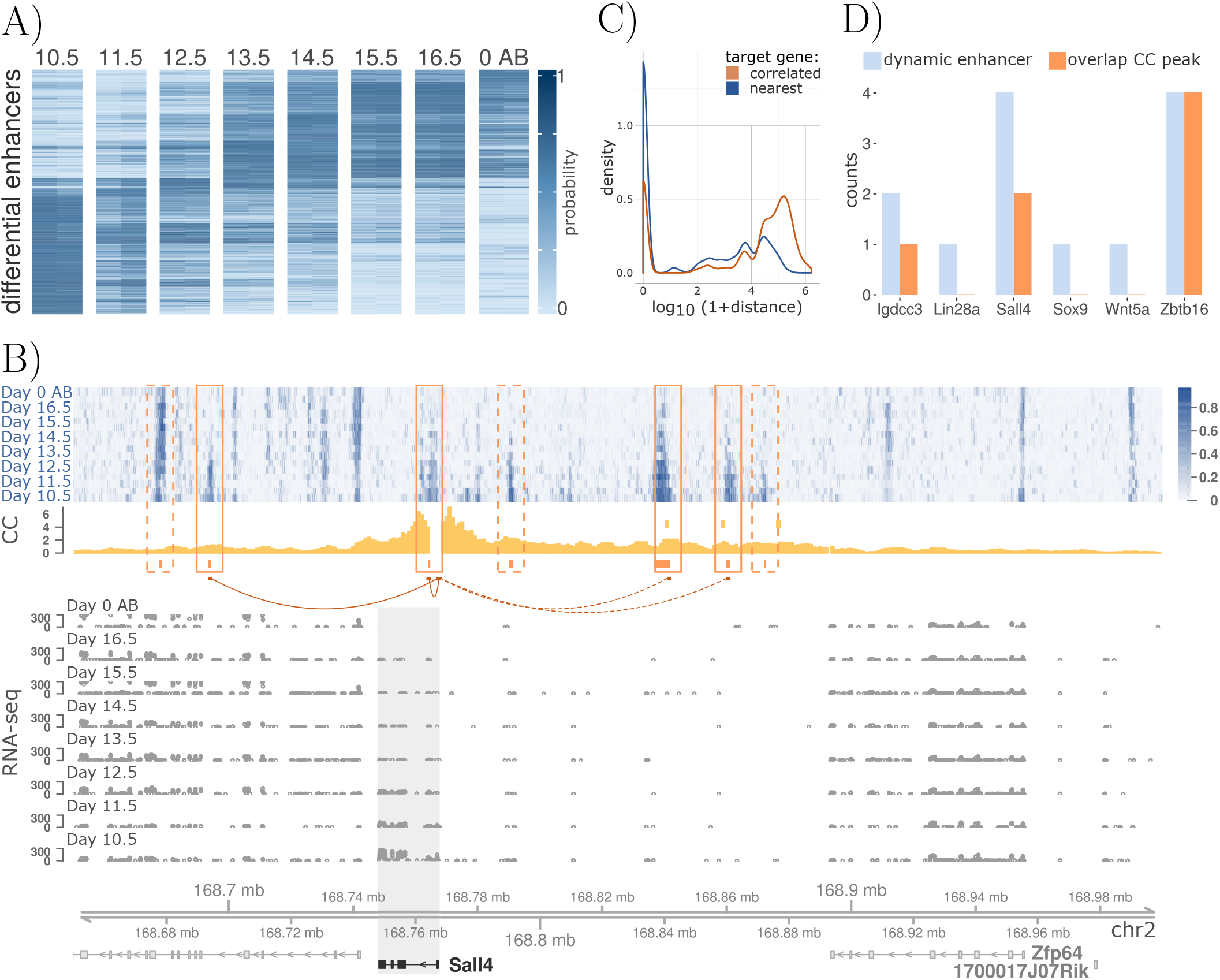
Differential enhancer-gene pairs in mouse embryo midbrain. **A)** Dynamic enhancer regions, colored by their respective enhancer probability, for eight time points (day 10.5 to day 0 after birth, *AB*) in mouse embryo midbrain. **B)** Enhancer probability tracks of the eight time points (blue) within a topologically associated domain. Eight differential enhancers could be assigned across all conditions (orange bars). Of these, enhancer probabilities of four regions (solid orange boxes) highly correlate with the gene expression of *Sall4* (bold black), a gene that regulates early embryonic development (orange arcs). CaptureC-seq data (*CC*) of mouse embryo midbrain (day 10.5) recapitulate these regulatory units (yellow histogram, yellow CC peak calls). **C)** Distribution of the log_10_ transformed distance between dynamic enhancers and their correlated target genes (orange) and the nearest gene (blue). **D)** Dynamic enhancers were filtered for regions that are active in mouse embryo midbrain at day 10.5. The number of differential enhancers found in the surrounding TAD of six genes that are active for mouse embryo midbrain at day 10.5 are displayed in light blue. The overlap with called CaptureC-seq peaks is shown in orange.

With this we build 258 regulatory units describing putative dependencies between differential enhancer regions and target genes located within the same TADs (see Section 5.6 for a description of the TAD calling software). Altogether, 61 of these differential enhancers are located within the gene body of the correlated target gene, whereas the remaining 197 regions are located in gene-free regions with a distance ranging from 75 bp to 1, 662, 424 bp to the correlated gene (see Figure 5 C). For about one third (64/197) of the intergenic regulatory units, the putative target gene is also the nearest gene to the respective differential enhancer region. Hence, the interaction range of a gene/promoter-enhancer pair is very heterogeneous and the nearest gene is not automatically the best choice for a target.

The majority of the regulatory units consists of single dynamic enhancer elements which are interacting with only one putative target gene (134/258). A small proportion of the regulatory units (55/258) rather describe genes that are correlated to multiple differential enhancers at once. Interestingly, several target genes seem to be regulated by the same enhancer region (35/258) which was also observed by van Arensbergen *et al.* (2014).

For four differentially active enhancers the probability patterns over all time points are highly correlated with the dynamic gene expression of *Sall4* (Figure 5 B), a known regulator in early embryonic development (Zhang *et al.*, 2006). We validate the results with CaptureC-seq experiments as exemplified by Andrey *et al.* (2017). Here we use interaction counts of mouse embryo midbrain CaptureC-seq (CC) data at day 10.5 (see Section 5.7 for details) with a viewpoint located at the promoter region of *Sall4*. Four differentially active enhancer regions are in close proximity to the three reported CC peaks and four additional regions could be only found with our **CRUP** framework. These additionally regions also show a slight increase in the interaction profile via visual inspection (yellow distribution, Figure 5 B). In total, we compared our predictions to Capture-C peaks for six different viewpoints located in the promoter regions of genes which are active at day 10.5 in mouse embryo midbrain and found an overlap with reported CC peaks for three of them (see Figure 5 D).

### 2.7 Regulatory units are well recapitulated by 3D chromatin structures

To further investigate the connection between predicted regulatory units and three-dimensional physical interactions between regulatory elements, we analyzed ultra-deep coverage Hi-C maps. We applied **CRUP** to a data set focusing on neural differentiation and cortical development in mice (Bonev *et al.*, 2017) comprising ChIP-seq, RNA-seq and Hi-C experiments across three developmental states: embryonic stem cells (ES), neural progenitor cells (NPC) and cortical neurons (CN).

We inferred 6, 855 regulatory units, which are clustered into 14 groups based on the activity pattern (see Section 5.9), and compared our results to log_2_ observed/expected (O/E) normalized Hi-C interaction matrices (see Section 5.6). Figure 6 shows a single regulatory unit, where the differential enhancer region is linked to the gene *Glul*. The Hi-C interaction frequencies across the three developmental states confirm the observed trend. The dynamic enhancer is predicted to be active only in ES cells but not in the other two conditions, which is also visible by normalized spatial interaction values. Next, we separately investigated clusters of regulatory units that are specific for only one condition. After dividing each interaction count triplet by its maximum value, the dynamic changes across the three conditions can be visualized for all regulatory units (Figure 6 B). These results not only confirm that cell type-specific gene-enhancer contacts are established concomitant with gene expression as already stated by Bonev *et al.* (2017), but they also show that dynamic enhancer activity goes hand in hand with physical changes in the three dimensional chromatin organization.

**Figure 6:**
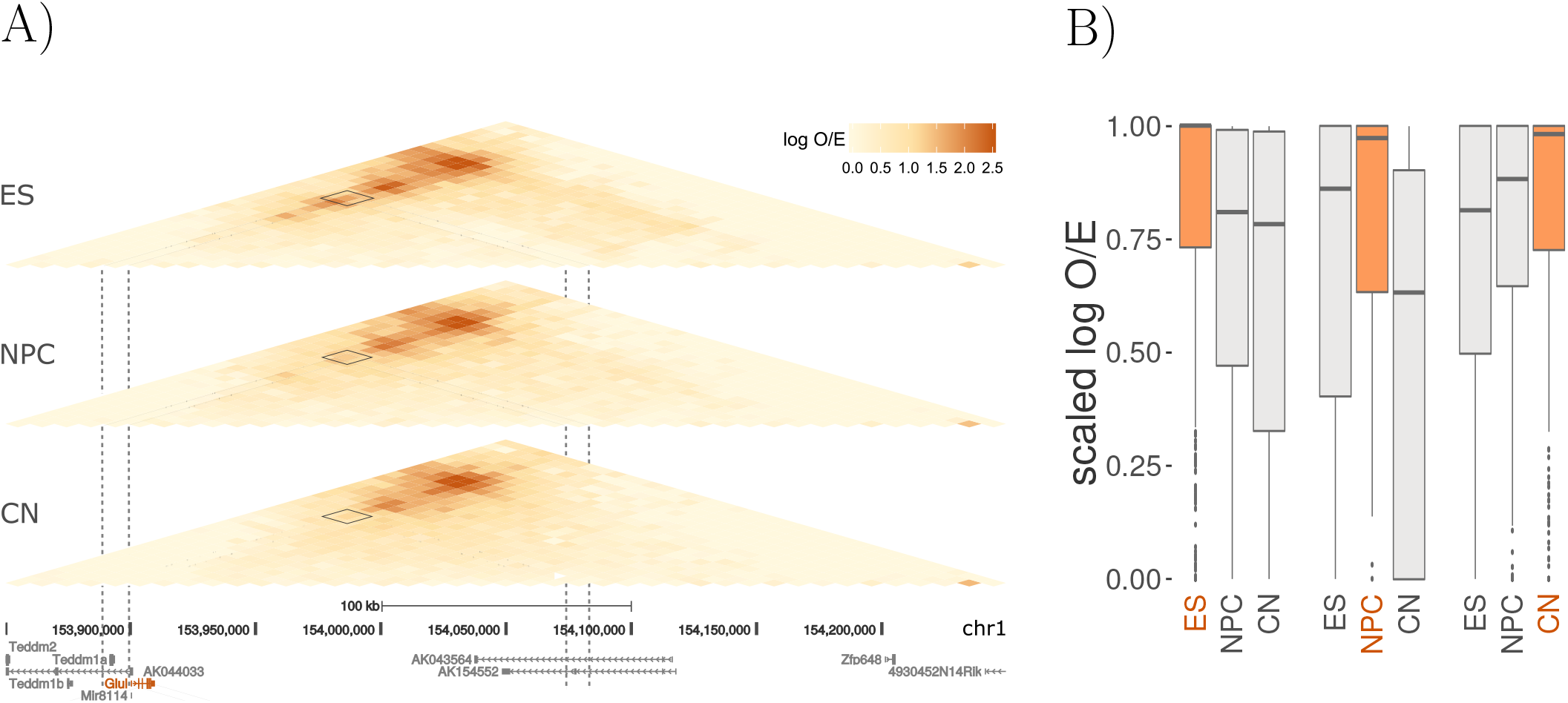
Differential regulatory units across mouse neural differentiation. **A)** Interaction matrices (log O/E) of three HiC-seq experiments of mouse embryonic stem cells (*ES*), neural progenitor cells (*NPC*) and cortical neurons (*CN*). A differential regulatory unit is indicated with dashed gray lines and a solid rectangle, showing the interaction of a differentially active enhancer region and the correlated gene *Glul* (red). **B)** Differentially active enhancers were filtered for regions that are only active in ES (left), only active in NPC (middle) and only active in CN (right), whereas the respective active condition is highlighted in orange. For these regions, normalized (log O/E) HiC interaction counts that overlap the predicted differential regulatory units were re-scaled to [0, 1], such that the highest interaction count for each region is 1.

### 2.8 Analysis of regulatory units in the context of a rheumatoid arthritis model

So far we evaluated our proposed framework **CRUP** to create condition-specific regulatory units on experiments focusing on developmental changes. Next, we apply our framework **CRUP** to a complex disease study which is part of the German Epigenome Program (DEEP, 2017), with the aim to identify the differences between two healthy mice and two mice which are affected by destructive rheumatoid arthritis (*’RA-like’*, see Section 5.1). Rheumatoid arthritis, an autoimmune inflammatory disease, affects approximately 0.5 − 1% of the human population and can lead to permanent joint destruction (Alamanos and Drosos, 2015).

We identified 514 differential enhancers and 462 differential regulatory units, of which about 60% (279) describe novel promoter/gene-enhancer pair activity that can only be found in the affected mice. We performed a motif analysis on all differential enhancer regions as described in Section 5.12. The TF motifs for KLF4, EGR2, IRF1 and FLI1 show higher enrichment in the cluster which contains enhancers that are solely active in the RA-like samples, and were already shown to be connected to rheumatoid athritis (Myouzen *et al.*, 2010; Luo *et al.*, 2016; Salem *et al.*, 2014; Sato *et al.*, 2014). A list of all enriched motifs is given in Figure S14.

Additionally, we performed a pathway analysis on all putative target genes that are correlated with differentially active enhancers in the affected mice using the Kyoto Encyclopedia of Genes and Genomes (KEGG) (Kanehisa and Goto, 2000; Kanehisa *et al.*, 2000, 2017), a curated database of molecular pathways and disease signatures (see Section 5.13 for details). Genes in the top five resulting KEGG pathways (see Table 1) have been previously associated with rheumatoid arthritis (Szekanecz *et al.*, 2010; Vogelpoel *et al.*, 2015; Chiffoleau, 2018; French and Yokoyama, 2004; Bergler-Czop *et al.*, 2016). A complete list of all gene-enhancer pairs and their associated top five KEGG pathway can be found in Table S6. A whole differential regulatory hub involving the CCR-gene cluster, which is part of the most significant pathway (Chemokine signaling pathway), is shown in Figure 7. Interestingly, the TF motif IRF1 is enriched in the differentially active enhancer which is correlated with all genes in the CCR-gene cluster. Interferon regulatory factor 1 (IRF1) is not only connected to rheumatoid athritis (Salem *et al.*, 2014) but is also a part of the C-type lectin receptor signaling pathway which we also found to be over-represented in out disease-related target gene list (see Table 1).

**Table 1:** KEGG pathway analysis results. Shown are the top five KEGG pathways over-represented in the putative target genes which are highly correlated with enhancer regions solely active in the samples with destructive arthritis (*’Genes’*). The list is sorted by the p-value for over-representation (*’N’* is the number of all genes in the respective pathways).

| Pathway ID | Pathway | N | #Genes | p-value |
| --- | --- | --- | --- | --- |
| path:mmu04062 | Chemokine signaling pathway | 27 | 13 | 1.655229e-06 |
| path:mmu04666 | Fc gamma R-mediated phagocytosis | 17 | 9 | 2.760813e-05 |
| path:mmu04625 | C-type lectin receptor signaling pathway | 23 | 10 | 8.401368e-05 |
| path:mmu04650 | Natural killer cell mediated cytotoxicity | 14 | 7 | 3.649052e-04 |
| path:mmu05167 | Kaposi sarcoma-associated herpesvirus infection | 28 | 10 | 5.757081e-04 |

**Figure 7:**
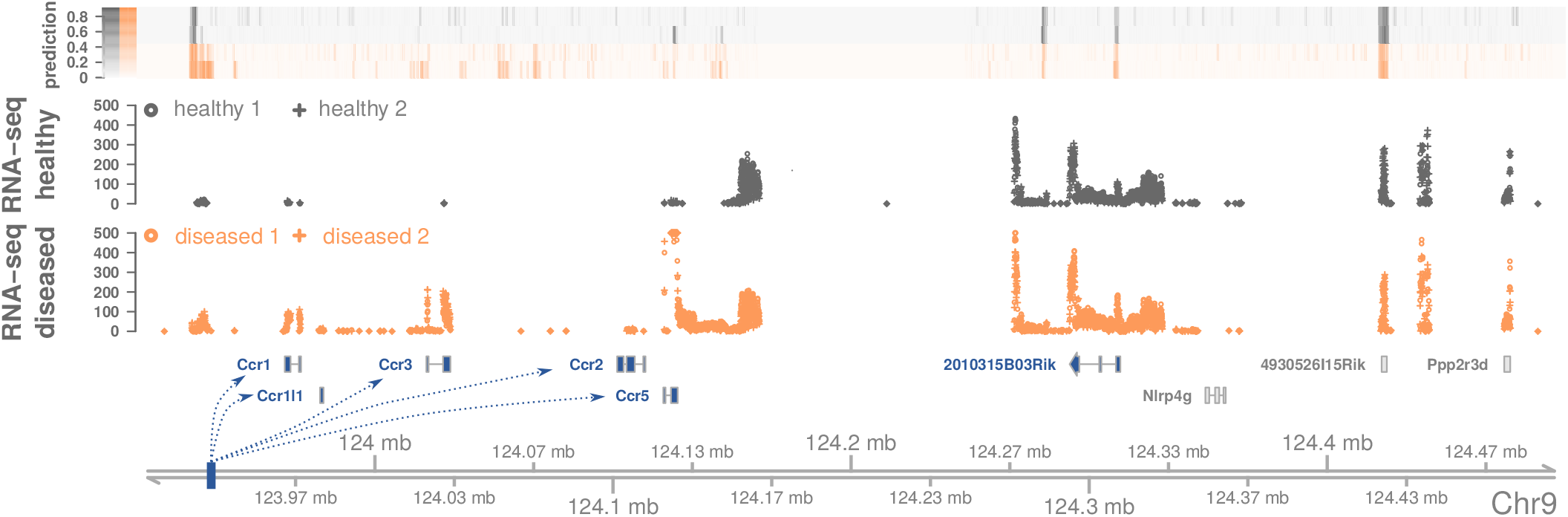
Example region containing differential regulatory units in the context of rheumatoid arthritis. Topologically associated domain and enhancer probabilities (’*prediction*’) of two healthy mice (gray) and two mouse model for destructive arthritis (orange). One enhancer in this region (blue bar) was found to be only active in the diseased samples. Using RNA-seq experiments of the same samples (displayed raw counts are cut at a maximum of 500), six genes are higly correlated (≥ 0.9, highlighted in blue) with the probabilities of the differential enhancer. Five of these genes (Ccr1, Ccr1l1, Ccr3, Ccr2, Ccr5) belong to the *CCR*-gene cluster that is an important part in the chemokine signaling pathway and also known to play a role in rheumatoid arthritis.

In summary, our framework **CRUP** is well suited to detect reliable candidate enhancer regions that act dynamically in different disease states as well as to link these enhancers to differentially expressed target genes building disease-associated regulatory units.

## 3 Discussion

In this work we described the three-step framework **CRUP** (**C**ondition-specific **R**egulatory **U**nits **P**rediction) to identify enhancer regions in a genome-wide manner, assign the predicted enhancers to different conditions and subsequently correlate the differential enhancers to putative target genes within their topologically associated domain to build condition-specific regulatory units.

We first showed that our random forest based enhancer classifier **CRUP-EP** can be reliably applied across different cell types and species without the need for re-training, solely depending on six core HMs. By integrating known structural characteristics of enhancer regions, namely an open region flanked by nucleosomes, into our feature modeling, our enhancer classifier accounts for the length of accessible chromatin and the spatial resolution of enhancer predictions. Our results show that the prediction performance of **CRUP-EP** across different cell types and species depends rather on the test than on the training data. We speculate that differences in ChIP-seq quality (see Figure S1) for certain training regions can be tolerated during the learning process and are not crucial for finding enhancer-specific HM pattern. However, for test regions, poor ChIP-seq signals very likely result in a decrease of performance. Another factor is the quality of the active enhancers which we defined based on the FANTOM5 database (see Table S3). While some weak or even mislabeled enhancers (false positives) in the training set still allow for a good enhancer representation by the classifier in terms of HM signals, mislabeled enhancers in the test set lead to false negatives predictions and thus directly reduce the recall results. Further, the highest number of suitable FANTOM5 cell lines for a confident enhancer definition was available for the mESC data set, which shows the best test set performance for almost all classifiers.

We further showed that our enhancer classification approach outperforms the unsupervised genome-segmentation tool ChromHMM and is comparable to another state-of-the-art random-forest based approach, REPTILE. In terms of transferability across different cell types and species, our classification approach even outperforms REPTILE. For this comparison, REPTILE was applied excluding differentially methylated regions (DMRs), which was part of the originally proposed feature set, as this was not available for all cell lines used in this study. Including DMRs led to an increase in spatial resolution which, nevertheless, remained below the results achieved with our classifier for the top 12, 500 predicted enhancers. Although the basic concept of the two supervised methods is similar, it is beneficial to include multiple windows in the feature set as we could show in our cross-validation, and to split the enhancer prediction into two separate tasks. This becomes especially visible when comparing the spatial resolution of the two classifiers based on the exact same training and feature setting. Another reason for the varying performance results across cell types/species could lie in the different normalization strategies. REPTILE does not offer an integrated normalization, but instead gives recommendations how to prepare the input data which we followed in our analysis. We advocate that a quantile normalization to the corresponding distribution of the data set used for training is crucial for a reliable enhancer prediction across cell types and species, and therefore incorporated this in our framework.

In a second step, **CRUP-ED**, we assign enhancers to different conditions using a permutation test on the enhancer probabilities obtained by the first module of **CRUP**. This approach can be applied to more than two conditions as the test is performed in a pair-wise manner. Using the resulting p-values we are able to create an activity pattern for each single bin which can then be used to combine and cluster all differentially active enhancers. We demonstrate that the assignment of clusters across different conditions is in good agreement with HM counts as well as with independent ATAC-seq data. Limitations arising from the raw data and from the enhancer prediction approach are consequently also reflected in the predicted differential enhancer regions. For instance, due to poor quality of individual HM ChIP-seq experiments the enhancer predictions might vary across samples in one condition and could therefore influence the results in the permutation test. Increasing the number of replicates could be one way to overcome this drawback since the implemented weighted difference between two conditions benefits from an enhanced sample size.

Lastly, we utilize **CRUP-ET** to integrate further genomic information, obtained from RNA-seq and Hi-C experiments, to link condition-specific enhancers to putative target genes via a *’greedy matching’* approach. To this end, we compute the correlation between normalized gene expression counts and enhancer probability values across all samples within the same TAD and put a strict threshold on the results to build high confidence regulatory units. Next, we evaluate our results by comparing regulatory units with Capture-C and Hi-C experiments. We could show that our inferred condition-specific gene-enhancer pairs are well recapitulated by physical dynamics in chromatin structures. To reduce the search space of interacting promoter/gene-enhancer pairs, we use TADs as a more sophisticated approach to form regulatory units rather than simply applying a distance based window. We show that the range in which differential enhancers and putative target genes are connected varies and that the nearest gene is often not the gene with the highest correlation. The resolution of Hi-C based experiments is still not on a single base pair level and might lead to wrongly associated promoter/gene-enhancer pairs, especially because the approach is also highly dependent on the performance of the TAD calling algorithm. We are utilizing TADs from murine stem cell experiments, to reduce the search space for detecting regulatory units for all the presented examples. We argue that these structures are highly stable across cell types and conserved in related species as observed in recent studies (Dixon *et al.*, 2012; Rao *et al.*, 2014). However, it was also shown, that structural differences between conditions occur, especially on a low scale sub-TAD level (Bonev *et al.*, 2017). Furthermore, the three dimensional landscape may change dramatically when structural variations disrupt the boundary structure as for example shown by Lupianez *et al.* (2015). In the future, condition specific Hi-C experiments could further help the presented approach in linking differentially active enhancers to putative target genes.

The complete framework was further applied to a complex disease study to identify differential regulatory units associated with rheumatoid arthritis. By applying motif analysis to the resulting differentially active enhancers, we were able to connect several regions to TF motifs that are linked to the disease. In combination with standard KEGG pathway analysis on the putative target genes we could show that our framework is well suited to identify candidate regulatory regions that behave differently depending on the disease state. To further validate these regions, additional follow-up experiments could complement the presented analysis.

The input to **CRUP** consists of a number of HM ChIP-seq experiments, each of which could in principle be analyzed by eye. Interpreting the combination of experimental tracks and, worse, many tracks under many conditions is, however, beyond the capacity of a human brain. As a result, many epigenetic experiments in the end get exploited only for studying the vicinity of a particular gene and do not serve the purpose of an unbiased, whole-genome inquiry. We thus see our method as an information integrator that reduces the diverse layers of information into an interpretable predictor, in turn allowing to rank signals across the entire genome.

## 4 Conclusion

In summary, we presented the three-step framework **CRUP, C**ondition-specific **R**egulatory **U**nits **P**redictions, to identify and assign differentially active enhancer regions in different states and link them to putative target genes within the same topologically associated domain.

The presented approach is user friendly as it aims to overcome the time consuming difficulties when comparing single read count tracks for several features and conditions. The framework is implemented in R and can be executed by solely providing mapped read counts for ChIP-seq and RNA-seq experiments.

Our pre-trained classifier can be used without the need of re-training and also outperforms existing methods especially when applied across various tissues and species. The resulting dynamically changing enhancer-gene pairs are in good agreement with three-dimensional interactions and can be used to further complement studies that aim to unravel dynamic epigenetic behaviour across different conditions.

## 5 Materials and Methods

### 5.1 Cell culture and isolation

#### Mouse embryonic stem cells

E14 mouse embryonic stem cells (mESCs) were cultured and routinely passaged every two days in ES medium plus leukemia inhibitory factor (LIF) in order to maintain the pluripotent state of the cells (Smith *et al.* 1988, Pease *et al.* 1990). To exit from pluripotency and push the cells towards differentiation, LIF was withdrawn and retinoic acid (RA) was added to the medium for a short pulse of 4h.

All experimental data related to these samples are accessible via GEO (GEO:GSE120376).

#### Mouse synovial fibroblasts

Murine SF (Synoial Fibroblasts) were isolated by enzymatic digestion from hind paws of 12 week old *hTNFtg* (reactive arthritis, strain Tg197 overexpressing human TNF) and wildtype (healthy control) as described before (Wehmeyer *et al.*, 2016; Keffer *et al.*, 1991).

#### Mouse adipocytes

Samples for adipocytes were isolated by collagenase treatment for 5 minutes followed by 5 minuts of collagenase inactivation as described before (Arrigoni *et al.*, 2016). After centrifugation the fat layer was collected.

#### Mouse hepatocytes

Primary mouse hepatocytes were obtained from two female mice (C57BL/6J x DBA/2 background) at the age of nine weeks. The isolation of primary mouse hepatocytes was performed by a two-step EDTA/collagenase perfusion technique as described by Godoy *et al.* (2013).

#### Human hepatocytes

Primary human hepatocytes were obtained from three different female donors (age 28-70 years) undergoing surgery due to primary or secondary liver tumors. Hepatocytes were isolated from healthy liver tissue remaining from liver resection as described in Godoy *et al.* (2013). Informed consent of the patients for the use of tissue for research purposes was obtained and experiments were approved by the local ethical committees.

### 5.2 Processing of histone modification ChIP-seq data

For all biological samples presented in this study, ChIP against six core HMs, H3K27ac, H3K27me3, H3K4me1, H3K4me3, H3K36me3 and H3K9me3, was performed. As a control served the sheared chromatin without antibody (Input). We utilized the tool plotFingerprint which is part of the deepTools project (Ramirez *et al.*, 2014) to assess quality metrics for all ChIP-seq experiments. Where we need to visualize read count enrichments in particular genomic regions, we employ the tool plotHeatmap which is also part of the deepTools project (Ramirez *et al.*, 2014).

#### Mouse embryonic stem cells

6 × 10^5^ low passage (< 10) E14 cells were cultivated for 48h in regular ES medium containing LIF. 4h prior to cross-link cells were treated with LIF or RA. Sequencing libraries were prepared and the resulting DNA fragments were paired-end 50bp sequenced on a Illumina HiSeq 2500 device. Raw sequencing reads were subsequently aligned to the genome assembly *’GRCm38’* with STAR (Dobin *et al.*, 2012) and duplicates where removed using Picard tools (Wysoker *et al.*, 2013).

#### Mouse synovial fibroblasts

ChIP-seq from 2 × 10^6^ cells was carried out as described before (Arrigoni *et al.*, 2016). Resulting DNA fragments were paired-end 50bp sequenced on a Illumina HiSeq 2500 device and raw sequencing reads were aligned to the genome assembly *’GRCm38’* using BWA-MEM (Li and Durbin, 2009; Li, 2013) and duplicates where removed using Picard tools (Wysoker *et al.*, 2013).

#### Mouse adipocytes

For mouse adipocytes chromatin from fixed cells has been extracted and sonicated for 15 minutes using Covaris S220 sonicator. Resulting DNA fragments were paired-end 50 bp sequenced on a Illumina HiSeq HiSeq 2500 device. Raw sequencing reads were aligned to the genome assembly *’GRCm38’* with BWA-MEM (Li and Durbin, 2009; Li, 2013) and duplicates where removed using Picard tools (Wysoker *et al.*, 2013).

#### Mouse hepatocytes

ChIP-seq was performed using 1 × 10^6^ primary mouse hepatocytes as was previously described (Kinkley *et al.*, 2016) with minor modifications. All six ChIP and input libraries from each sample were then pooled and paired-end sequenced on an HiSeq 2500 device. Raw sequencing reads were aligned to the genome assembly *’GRCm38’* with STAR (Dobin *et al.*, 2012) and duplicates where removed using Picard tools (Wysoker *et al.*, 2013).

#### Human hepatocytes

ChIP-seq was performed using 1 × 10^6^ primary human hepatocytes as was previously described (Kinkley *et al.*, 2016) with minor modifications. All six ChIP and input libraries from each sample were then pooled and paired-end sequenced on an HiSeq 2500 device. Raw sequencing reads were aligned to the genome assembly *’hs37d5’* with BWA-MEM (Li and Durbin, 2009; Li, 2013) and duplicates where removed using Picard tools (Wysoker *et al.*, 2013).

#### Mouse embryo midbrain

Raw reads from ChIP-seq experiments were downloaded from GEO (GEO:GSE88517, Gorkin *et al.* (2017)) and aligned to the genome assembly *’GRCm38’* with BWA-MEM (Li and Durbin, 2009; Li, 2013) and duplicates where removed using Picard tools (Wysoker *et al.*, 2013).

#### Samples in the context of mouse neural differentiation

Raw data from RNA-seq for the three in vitro generated murine cell types ES, NPC and CN were downloaded via GEO (GEO:GSE96107, Bonev *et al.* (2017)) and aligned to the genome assembly *’GRCm38’* with BWA-MEM (Li and Durbin, 2009; Li, 2013) and duplicates where removed using Picard tools (Wysoker *et al.*, 2013).

### 5.3 Processing of RNA-seq experiments

#### Mouse embryonic stem cells

2 × 10^5^ low passage (< 10) E14 cells were plated and cultivated for 48h in regular ES medium containing LIF. 4h prior to harvest, medium was exchanged and cells were treated with LIF or RA. Cells were harvested and three biological triplicates were subjected to RNA extraction. Sequencing libraries were generated from total mRNA input and high-throughput sequencing was performed on an Illumina HiSeq 2500 device generating resulting in 50bp paired-end reads. Raw reads were subsequently mapped to the mouse genome build *’GRCm38’* using BWA-MEM (Li and Durbin, 2009; Li, 2013).

#### Mouse synovial fibroblasts

Long RNA libraries were prepared from total mRNA input and sequenced on an Illumina HiSeq 2500 device resulting in 50bp and 100bp long paired-end reads. Raw reads were subsequently mapped with TopHat2 (Kim *et al.*, 2013) to the mouse genome build *’GRCm38’*.

#### Mouse adipocytes

RNA isolation for cells was performed using 1 ml TRIzol per sample followed by Isopropyl alcohol/Ethanol precipitation. Sequencing libraries were generated from total mRNA input and high-throughput sequencing was performed on an Illumina HiSeq 2500 device generating resulting in 100bp paired-end reads. Raw reads were mapped with TopHat2 (Kim *et al.*, 2013) to the mouse genome build *’GRCm38’*

#### Mouse hepatocytes

RNA was extracted from ~ 5 × 10^6^ hepatocytes homogenized in 1 mL Trizol. Sequencing libraries were generated from total mRNA input using TruSeq v3 Kit (Illumina) according to manufacturers instructions and high-throughput sequencing was performed on an Illumina HiSeq 2500 device generating resulting in 100bp paired-end reads. Raw reads were mapped to the mouse genome build *’GRCm38’* using BWA-MEM (Li and Durbin, 2009; Li, 2013).

#### Human hepatocytes

RNA was extracted from ~ 5 × 10^6^ hepatocytes homogenized in 1 mL Trizol. Sequencing libraries were generated from total mRNA input using TruSeq v3 Kit (Illumina) according to manufacturers instructions and high-throughput sequencing was performed on an Illumina HiSeq 2500 device generating resulting in 100bp paired-end reads. Raw reads were mapped with TopHat2 (Kim *et al.*, 2013) to the genome build *’hs37d5’*.

#### Mouse embryo midbrain

Raw reads from RNA-seq experiments were downloaded from GEO (GEO:GSE88517, Gorkin *et al.* (2017)) and aligned to the genome assembly *’GRCm38’* with STAR (Dobin *et al.*, 2012).

#### Samples in the context of mouse neural differentiation

Raw data from RNA-seq for the three in vitro generated murine cell types ES, NPC and CN were downloaded via GEO (GEO:GSE96107, Bonev *et al.* (2017)) and aligned to the genome assembly *’GRCm38’* with BWA-MEM (Li and Durbin, 2009; Li, 2013).

### 5.4 Processing of DNase-seq experiments

To compare open chromatin sites to HM signals, read counts from DNase-seq experiments were summarized for adjacent 100bp bins using the R package bamProfile (Mammana and Helmuth, 2016). Read count enrichments are visualized with the plotHeatmap funciton implemented in the software package deepTools (Ramirez *et al.*, 2014).

#### Mouse embryonic stem cells

Raw reads from DNase-seq experiments from mESCs (E14, Embryonic day 0) were downloaded from GEO (accession Nr.:GSM1014154) and aligned to the genome assembly *’GRCm38’* with BWA-MEM (Li and Durbin, 2009; Li, 2013). Duplicates were further removed using Picard tools (Wysoker *et al.*, 2013).

#### Mouse synovial fibroblasts

5 − 7 × 10^6^ nuclei were digested with DNaseI in 5 different dilutions as described before (Schmidt *et al.*, 2016). Raw sequencing reads were aligned to the genome assembly *’GRCm38’* with BWAMEM (Li and Durbin, 2009; Li, 2013) and duplicates where removed using Picard tools (Wysoker *et al.*, 2013).

#### Mouse adipocytes

Nuclei extracted from ~ 10 × 10^6^ nuclei by treatment with IGEPAL were digested with different concentrations of DNaseI as described before (Schmidt *et al.*, 2016) and kept at 4°C until further processing. Sequencing libraries were prepared and sequenced on an Illumina HiSeq 2500 device resulting in 100bp long paired-end reads. Raw sequencing reads were aligned to the genome assembly *’GRCm38’* with BWA-MEM (Li and Durbin, 2009; Li, 2013) and duplicates where removed using Picard tools (Wysoker *et al.*, 2013).

#### Mouse hepatocytes

Nuclei extracted from ~ 10 × 10^6^ nuclei by treatment with IGEPAL were digested with different concentrations of DNaseI as described before (Schmidt *et al.*, 2016) and kept at 4°C until further processing. Sequencing libraries were prepared and sequenced on an Illumina HiSeq 2500 device resulting in 100bp long paired-end reads. Raw sequencing reads were aligned to the genome assembly *’GRCm38’* with BWA-MEM (Li and Durbin, 2009; Li, 2013) and duplicates where removed using Picard tools (Wysoker *et al.*, 2013).

#### Human hepatocytes

Nuclei extracted from ~ 10 × 10^6^ nuclei by treatment with IGEPAL were digested with different concentrations of DNaseI as described before (Schmidt *et al.*, 2016) and kept at 4°C until further processing. Sequencing libraries were prepared and sequenced on an Illumina HiSeq 2500 device resulting in 100bp long paired-end reads. Raw sequencing reads were aligned to the genome assembly *’hs37d5’* with BWA-MEM (Li and Durbin, 2009; Li, 2013) and duplicates where removed using Picard tools (Wysoker *et al.*, 2013).

### 5.5 Processing of ATAC-seq experiments from mESC

2×10^5^ low passage (< 10) E14 cells were cultivated for 48h in regular ES medium containing LIF. 4h prior to harvest, cells were treated with LIF or RA (1 M). 75000 cells per treatment were subjected to transposition reaction and PCR amplification of accessible regions by Omni-ATAC-seq as described previously by Corces *et al.* (2017). Sequencing libraries were constructed and DNA fragments were paired-end 50bp sequenced on a Illumina *HiSeq 4000* device. Raw reads were subsequently aligned to the mouse genome build GRCm38m using BWA-MEM (Li and Durbin, 2009; Li, 2013) and duplicates were removed upon filtering using SAMtools (Li *et al.*, 2009). *ATAC-seq* peaks were idenitfied using MACS2 (Zhang *et al.*, 2008).

### 5.6 Processing of HiC-seq experiments

The Juicertools command ’*dump*’ (Durand *et al.*, 2016) was used to extract data from Hi-C archives associated with three in vitro generated murine cell types ES, NPC and CN (Bonev *et al.*, 2017):

- http://hicfiles.s3.amazonaws.com/external/bonev/ESmapq30.hic
- http://hicfiles.s3.amazonaws.com/external/bonev/NPCmapq30.hic
- http://hicfiles.s3.amazonaws.com/external/bonev/CNmapq30.hic

With this each matrix is Knight-Ruiz (KR) normalized (Knight and Ruiz, 2013) at 10*kb* resolution and the observed/expected (O/E) ratio is computed. For visualization O/E interaction maps were further log_2_ converted and negative values were set to 0. Additionally, topologically associated domains (TADs) were identified by utilizing TopDom (Shin *et al.*, 2016) on 25*kb* binned and KR normalized matrix based on murine stem cells (ES) using a window of 750 kb (30 × 25kb) for the TopDom algorithm. These regions were used to reduce the search space for promoter/gene-enhancer interactions.

### 5.7 Filtering of Capture-C experiments for mouse embryo midbrain

*Capture-C-seq* profiles from mouse embryo midbrain (Day 10.5) were downloaded from GEO (GEO:GSE84795, Andrey *et al.* (2017)) and coordinates were transferred to the mouse genome build *’GRCm38’* utilizing the function *liftOver* which is implemented in the R package *rtracklayer*. We further filtered the provided list of chromatin states that were assigned to each promoter by EpiCseq (Mammana and Chung, 2015) for the state’*Active A*’, which resembles active promoters. The result was further filtered for genes present in any differential regulatory unit that was found across all conditions in mouse embryo midbrain (Gorkin *et al.*, 2017), resulting in six genes.

### 5.8 CRUP-EP: Enhancer Prediction

#### Preparation and normalization of HM counts

Histone modification count signals are summarized for adjacent non-overlapping 100 bp bins utilizing the R package *bamProfile* (Mammana and Helmuth, 2016), following a log_2_ input normalization (with pseudo count of 1) of the raw counts. We compute the log_2_ ratio (also with pseudo count of 1) between H3K4me1 and H3K4me3 after shifting the distribution of their input-normalized count values to ≥ 0.

Before making predictions on a sample with our classifier, the input-normalized count values are quantile normalized to the corresponding distributions of the data used for training. This is done with the *normalize.quantiles.target* function of the R package *preprocessCore* (Bolstad, 2018).

#### Definition of high confidence enhancer regions

One specific hallmark for enhancer activity was found to be the initiation of RNAPII transcription, which was used by the FANTOM5 project (Andersson *et al.*, 2014). Short RNA-seq and CAGE was applied to a variety of different cell types and tissues to detect bidirectional capped transcripts. CAGE count data were downloaded for mouse adipocyte cells, mouse embryonic stem cells, mouse fibroblast cells as well as human and mouse liver cells. Depending on the number of available replicates for each cell line we chose different cutoffs for the CAGE counts to define a first set of putative enhancers according to the summary in Table S3. To get our final high-confidence enhancer set we centered the putative FANTOM5 enhancers based on DNase-seq peaks and discarded enhancers without any overlap with DNase-seq peaks as summarized in Table S4. To convert the genome coordinates of the enhancer regions given by the FANTOM5 project from genome build *GRCm37* to *GRCm38* we applied the Batch Coordinate Conversion tool *liftOver* from the UCSC Genome Browser Utilities (Hinrichs *et al.*, 2006).

#### Definition of active and inactive promoter regions

For murine ESC, adipocytes, liver and fibroblast cells, and for human liver cells we computed FPKM gene expression values from RNA-seq data as described in Section 5.3.

Based on the gene annotations from the Ensembl data base (GRCh37.70 and GRCm38.90), we defined a gene with an FPKM value greater than two as active and a gene with FPKM value of zero as inactive (0 < FPKM <= 1 was not used for training). In case replicates were available, all of the replicates had to fulfill the chosen FPKM cutoff to be accounted to the one or the other class. An exemplary distribution of FPKM values, here for mESC^+^, can be seen in Figure S4. Building up on this, we then defined an inactive promoter as the 100 bp bin overlapping the TSS of an inactive gene. An active promoter is defined as the 100 bp bin having an overlap with the TSS of an active gene as well as with a DNase-seq peak in the corresponding cell type. An overview can be found in Table S5.

#### Enhancer prediction based on random forests

We use a combination of two binary random forest classifier for our enhancer prediction, where both consist of *M* = 70 decision trees. The first classifier (classifier 1) learns the difference between active genomic regions (active promoters, enhancers) and inactive genomic regions (inactive promoters, remaining intra- and intergenic regions). The second one (classifier 2) learns to distinguish enhancers from active promoters, such that it gives the probability of a region to be an enhancer assuming it is an active region. The final enhancer probability assigned to each 100bp bin, bin_*x*_, in the fragmented genome is the product of both classifiers according to Bayes’ theorem:

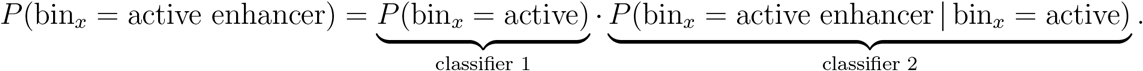

In the two distinct training sets for classifier 1 and 2 we emulate a typical genome composition as reported, e.g., in Kellis *et al.* (2014). The training set of classifier 1 is composed of 10% enhancers, 2% active promoters, 2% inactive promoters, 6% intragenic and 80% intergenic regions, summing up to 1000 regions in total. Classifier 2 is trained on 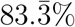 enhancers and 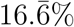 active promoters. Here we keep the same enhancer/promoter ratio and total numbers than in the first training set, i.e., we always use 120 regions selected according to these rules. Overall, this also serves the purpose of adequately reflecting the imbalance between enhancers and non-enhancer regions in the genome. The feature set, which is also chosen individually for the two classifiers, is derived from summed and normalized ChIP-seq read counts for the six core HMs. For classifier 1, we consider only H3K27ac, H3K27me3 and H3K9me3, whereas for classifier 2 we consider all six core HMs as well as the H3K4me1/me3 ratio.

Since we want to represent the physical structure of an enhancer (nucleosome - accessible region -nucleosome) we divide a large window of 1100 bps into 11 non-overlapping bins, i.e., the center bin (bin_*x*_) plus *N* = 5 bins on either side, resulting in a total number of 11 · 3 = 33 features for classifier 1 and 11 · 7 = 77 features for classifier 2.

The number of neighboring bins *N* in the feature set, as well as the number of decision trees *M* in the random forest are parameters that we optimized according to the description in the following section.

#### Parameter tuning

We used 5-fold cross-validation over 10 different training seeds to find the optimal number of decision trees *M* ∈ {10, 20, …, 100} and neighboring windows *N* ∈ {0, 1, …, 10}. Each of the 10 training sets used is chosen as described in the previous paragraph. Based on the AUC-PR (area under the PR curve) performances (see Figure S12), we fixed the combination of *N* = 5 neighboring windows and *M* = 70 trees for both classifiers. With the optimized parameter choice we trained classifier 1 and 2 on two final randomly sampled training sets which can have a possible overlap with the 10 training sets used for parameter tuning. The parameter setting of *N* = 5 and *M* = 70 is used in all our analyses.

#### Enhancer peak calling and building of enhancer clusters

Genome-wide predictions result in enhancer probability values for each 100 bp bin in the genome which are further summarized to define enhancer peaks. To this end, all bins with a probability ≥ 0.5 are sorted in descending order according to their probability value and expanded by five bins up and downstream resulting in a window length of 1100 bps. By going through the sorted list of high probability regions, starting with the highest probability, all windows that overlap the current window are discarded. This results in a sorted list of non-overlapping enhancer peaks of length 1100 bp.

Enhancer peaks are further summarized into enhancer clusters solely considering the distance between them (maximum distance of 12.5 kb), which partly reflects the definition of super-enhancers as stated by Whyte *et al.* (2013) and Love *et al.* (2013).

### 5.9 CRUP-ED: Enhancer Dynamics

#### Statistical inference of differences between two conditions

Enhancer probabilities for all 100 bp bins and samples are collected in a matrix *A* = (*A*_*xi*_) where *A*_*xi*_ corresponds to bin_*x*_ in sample *i*. In the following we denote by *A*_*C*^1^_ = (*A*_*xi*_)_*i*∈*C*^1^_ the sub-matrix of *A* with columns corresponding to samples from condition *C*^1^ (applies equally for condition *C*^2^). As the number of samples in each group is usually very small, we perform a non-parametric permutation test on the data set to compute an empirical distribution. This approach was already introduced in earlier studies, as for example by Tusher *et al.* (2001). First, all enhancer probabilities *A*_*xi*_ are independently shuffled for each sample *i*. The t-test statistic *T*_*x*_ is then calculated for each bin_*x*_ to obtain the weighted difference between the two conditions:

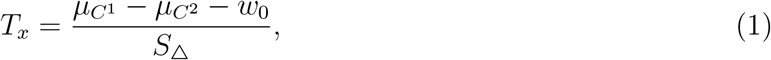

whereas *μ*_*C*^1^_ = *μ*(*A*_*xC*^1^_) and *μ*_*C*^2^_ = *μ*(*A*_*xC*^2^_) are the respective group means for bin_*x*_ and *w*_0_ defines the minimum difference between them (here: *w*_0_ = 0.5). *S*_Δ_ is the pooled standard deviation based on the group variances 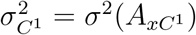 and 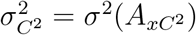:

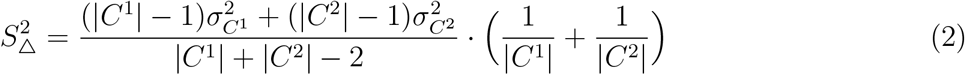

Empirical p-values for each bin_*x*_, *P*_*x*_ = *P*_*x*_(*C*^1^, *C*^2^), are obtained by counting the values *T*_*x*_ in the sampling distribution that exceed the true weighted difference 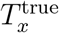, which means that the lowest possible p-value is 1/(1 + length of genome). By setting a cut-off *P*\* (default: 0.01) to the obtained *P*_*x*_ the genome is reduced to high confidence enhancer regions of length 100 bp that significantly differ in probabilities between two distinguishable conditions. Note that *S*_Δ_ is set to a small number ≈ 0 if |*C*^1^| = 1 and |*C*^2^| = 1 to avoid division by zero.

#### Clustering of differential enhancers using ‘activity pattern’

Significant differential enhancer regions of length 100 bp are obtained for all pairwise comparisons between any two conditions {*C*_1_, *C*_2_} ∈ *C* as described in the previous paragraph.

n the following the indicator function *T*(*C*^1^,*C*^2^) = *T_x_*(*C*^1^,*C*^2^) denotes if binx is an active enhancer in condition *C*_1_ but not in condition *C*_2_:

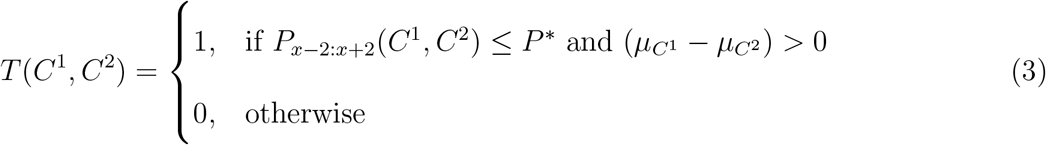

Note that additional to the p-value assigned to bin_*x*_, the p-values of two additional bins up and downstream of bin_*x*_ are required to be smaller than *P*\*. In the following bin_*x*_ is renamed as 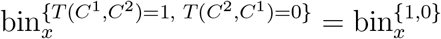 if the empirical p-values *P*_*x*−2:*x*+2_(*C*^1^, *C*^2^) ≤ *P*\* and if the difference in the group means (*μ*_*C*^1^_ − *μ*_*C*^2^_) > 0. The region will be denoted as 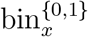 if *T*(*C*^2^, *C*^1^) = 1 and as 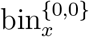 if *P*_*x*−2:*x*+2_(*C*^1^,*C*^2^) > *P*\*. With this, each differential enhancer bin_*x*_ can be allocated to a unique *’activity pattern’*, either {1, 0}, {0, 1} or {0, 0} (see Figure 8 for an overview). This notation expands as the number of conditions, |*C*|, increases. For example, if |*C*| = 3, the number of possible comparisons is 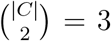, namely (*C*^1^, *C*^2^), (*C*^1^, *C*^3^) and (*C*^2^, *C*^3^). As each tupel can be assigned to three activity pattern, the total number of possible outcomes sums up to 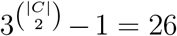, whereas the pattern {0, 0, 0, 0, 0, 0} does not include any differential information and can be discarded from the list.

**Figure 8:**
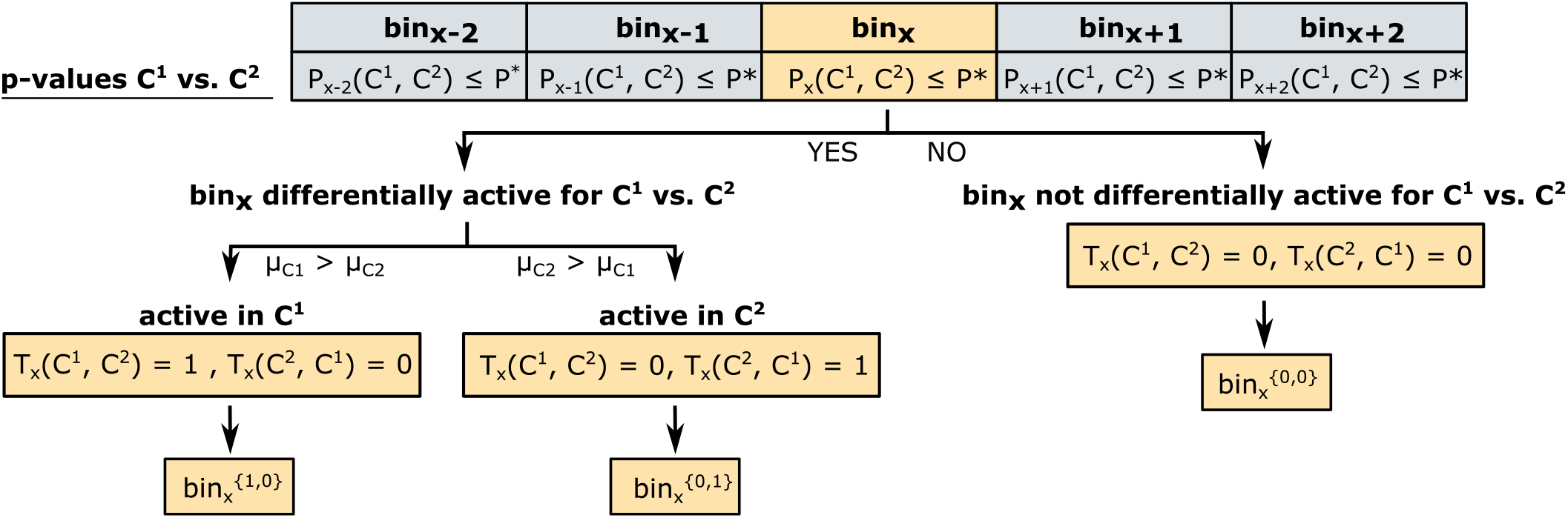
Assignment of activity pattern for the comparison of two conditions. For the differential comparison of enhancers in two conditions, *C*^1^ and *C*^2^, the activity pattern assigned to bin_*x*_ depends on the empirical p-values of bin_*x*_ and two neighboring bins to both sides (bin_*x*−1_, bin_*x*−2_, bin_*x*+1_, bin_*x*+2_). If one of the five empirical p-values exceeds the cutoff *P*\*, bin_*x*_ does not represent a differential enhancer between *C*^1^ and *C*^2^, and is assigned the activity pattern {*T*_*x*_(*C*^1^, *C*^2^) = 0, *T*_*x*_(*C*^2^, *C*^1^) = 0} = {0, 0}. If all five bins show an empirical p-value below *P*\* and the group mean of *C*^1^ is greater than the group mean of *C*^2^ (*μ*_*C*^1^_ *> μ*_*C*^2^_, bin_*x*_ represents an active enhancer in *C*^1^ and is assigned the activity pattern {*T*_*x*_(*C*^1^, *C*^2^) = 1, *T*_*x*_(*C*^2^, *C*^1^) = 0} = {1, 0}. In the opposite case (*μ*_*C*^2^_ > _*μ*_*C*1), bin_*x*_ is active in *C*^2^ with an activity pattern of {*T*_*x*_(*C*^1^, *C*^2^) = 0, *T*_*x*_(*C*^2^, *C*^1^) = 1} = {0, 1}.

The total range of all bin_*x*_ that are associated with the same activity pattern is summarized within a 2 kb distance whereas the bin_*x*_ with the lowest p-value *P*_*x*_ is stored as peak. If regions with different activity patterns are overlapping, these are combined and labeled with the activity pattern according to the lowest peak p-value.

### 5.10 CRUP-ET: Enhancer Targets

#### Regulatory units by ‘greedy matching’

Differential enhancer regions for any set of conditions *C* are obtained as described and clustered as described in Section 5.9. Gene expression counts per exon are obtained from RNA-seq experiments of the same conditions (see Section 5.3) using the function *summarizeOverlaps* implemented in the R package *GenomicAlignments* (Lawrence *et al.* (2013), v1.14.2). Summarized counts per gene are variance stabilized across the mean using the function *vst* implemented in the R package *DESeq2* (Love *et al.* (2014), v1.18.1).

All genes and differential enhancer regions are gathered within the same topologically associated domain (see Section 5.6). To find regulatory units of gene-enhancer pairs that behave similarly across conditions we apply a ‘greedy matching’ strategy. For this, Pearson correlation values are calculated between enhancer probability values and normalized gene expression counts across all conditions. All enhancer-gene pairs with a correlation ≥ 0.9 are considered as putative regulatory units and are reported.

### 5.11 Comparison to other enhancer predicting methods

#### Application of ChromHMM

ChromHMM (Kellis *et al.*, 2014) was applied to six core HMs to generate three genome-wide segmentations for undifferentiated mESCs based on *K* ∈ {8, 12, 16} chromatin states (Figure S8). For *K* = 8 we were not able to clearly separate an enhancer from the promoter state. For *K* = 12 and *K* = 16 we defined enhancers based on combinations of states with high emission probabilities for the enhancer marks H3K4me1 and H3K27ac, low emission probabilities for the repressive marks H3K27me3 and H3K9me3, and also low emission probabilities for the promoter mark H3K4me3. We tested four different enhancer definitions for *K* = 12 including states (i) *E*3, (ii) *E*3 + *E*12, (iii) *E*2 + *E*3 and (iv) *E*2 + *E*3 + *E*12, and for *K* = 16 the enhancer definitions are composed of states (i) *E*1, (ii) *E*1 + *E*16, (iii) *E*1 + *E*6 and (iv) *E*1 + *E*6 + *E*16.

The prediction performances of the defined enhancer state (versus all other states) for *K* = 12 and *K* = 16 were calculated based on the same ten test sets generated through different random seeds as used in Section 2.2. To determine an overlap we extend our test regions to 1100 bps centered on the respective region. Based on these definitions the numbers of true and false positives and negatives could be calculated.

#### Application of REPTILE

REPTILE (He *et al.*, 2017) was trained on different mouse (ESC, fibroblasts, adipocytes, liver) and human (liver) data. We first RPM normalized the ChIP-seq tracks and then performed a log_2_ input normalization on all HM data as recommended in the REPTILE paper.

For mESC, we made genome-wide predictions whereas for the other samples we only predicted on a test set. To do so, we chose the training set for REPTILE similarly as for our method (see 5.8), i.e. also trying to emulate a typical genome composition.

Genome wide predictions on mESC were generated using four different training and feature set combinations:

i. FANTOM5 derived enhancers and six core HMs
ii. p300 defined enhancers, six core HMs and intensity deviation
iii. p300 defined enhancers and ENCODE HMs
iv. p300 defined enhancers, ENCODE HMs and differentially methylated regions (DMRs)

Here, the six core HMs are from the in-house mESC data, the ENCODE HM data and the differentially methylated regions (DMRs) are taken from He *et al.* (2017). The intensity deviation for a specific target sample is described in He *et al.* (2017) as the signal/intensity of the target sample subtracted by its mean intensity in reference samples. In our setting, we included additional to the mESC target sample also the intensity deviation between intensity from mESC and the 13 data sets from our test set prediction across different tissues (Section 2.3).

Using the REPTILE peak calling tool with a probability threshold of 0.5 for the different scenarios, we got (i) 24,823, (ii) 34,584, (iii) 32,797 and (iv) 30,360 annotated enhancer regions.

### 5.12 Motif enrichment analysis

We performed motif hit enrichment analyses with the R package *motifcounter* (Kopp and Vingron, 2017) on individual enhancers or clusters of enhancers. The method is based on a higher-order Markov background model to compute the expected motif occurrences (hits) and a compound Poisson approximation for enrichment testing. We use the default parameters for the order of the background model and the false positive level for motif hits, order = 1 and *α* = 0.001, respectively. In our analysis of enhancer clusters, we refer to the fold-enrichment value for the over-representation of a motif. For a single enhancer sequence, we filter motifs by p-value (≤ 0.05) and individual motif hits by score (maximum) to pinpoint relevant TFBSs.

We tested for enrichment of the binding profiles of 579 TFs in total which were downloaded from the non-redundant JASPAR 2018 CORE vertebrate collection (Khan *et al.*, 2018) of position frequency matrices (PFMs).

### 5.13 KEGG pathway analysis

We used the curated database of molecular pathways and disease signatures to perform a over-representation analysis for KEGG (Kyoto Encyclopedia of Genes and Genomes) pathways (Kanehisa and Goto, 2000; Kanehisa *et al.*, 2017, 2000). To this end we applied the function *kegga* (*’species.KEGG = “mmu”, trend = T’*) implemented in the *edgeR* R package (McCarthy *et al.*, 2012; Robinson *et al.*, 2010) to identify murine KEGG pathways that are over represented in putative target genes that were found to be highly correlated with enhancer regions that are solely active in mice with rheumatoid athritis (correlation ≥ 0.9). As background we used all genes (R package *’Txdb.Mmusculus.UCSC.mm10.knownGene’*, Team and Maintainer (2016)) that are located within the same TADs as all identified regulatory units. We used the p-value (*’P.DE’*) to order the results and reported the best five pathways.

## Declarations

### Ethics approval and consent to participate

Not applicable.

### Consent for publication

Not applicable.

### Availability of data and materials

Data for ChIP-seq, RNA-seq, DNase-seq and ATAC-seq experiments for the pluripotent (mESC^+^) and differentiated (mESC^−^) mouse embryonic stem cell samples are available via GEO (GSE120376).

### Availability of the Application

The code to run the framework CRUP is available at https://github.com/VerenaHeinrich/CRUP.

### Authors’ contributions

A.R. and V.H. implemented the framework, did the analyses and wrote the manuscript. M.V. conceptualized the work and edited the manuscript. P.B. and R.S. helped with the analysis. L.V.G was responsible for mESC culture work and performed RNA- and ATAC-seq experiments, which was supervised by S.H.M., A.F. performed ChIP-seq experiments on mESC^+/−^ cells. T.P. initiated and directed the rheumatoid arthritis experiments and contributed to the editing of the manuscript. C.C. and J.H. provided human hepatocytes. A.K. initiated and directed mouse hepatocytes experiments. A.P. initiated and directed mouse adipocytes experiments. N.L. and S.K. performed animal handling and experiments of mouse hepatocytes and mouse fibroblast experiments. A.H. performed the animal handling and cell culture experiments of the rheumatoid arthritis models (mouse fibroblast) and contributed to the editing of the manuscript. B.C. and S.M.K. performed animal handling and experiments of human adipocytes and mouse hepatocytes experiments. J.L. performed animal handling and prepared the mouse hepatocytes samples and S.H. performed the data handling and analysis. S.K. and N.L. performed ChiP-seq of sequencing of mouse hepatocytes and human hepatocytes. A.L. performed ChIP-seq experiments on human hepatocytes and mouse adipocytes, which was supervised by T.M., N.G. performed Dnase-seq for all DEEP related experiments, which was supervised by J.W., X.Y. did basic analysis of mouse hepatocytes, which was supervised by H.C., S.H. did basic analysis of mouse adipocytes.

## Acknowledgements

We thank Andreas Richter and Karl Nordström for data handling and transfer. Many thanks to Edgar Steiger, Tobias Zehnder, Giuseppe Gallone and Jaydeep Bhat for their valuable comments and inspiring discussions. We also acknowledge Gilles Gasparoni for sequencing of DEEP DNAse-samples and the Deep-Sequencing Unit of the MPIIE (Emily Betancourt, Ulrike Boenisch and other technical staff). Finally, we also would like to thank all members of the DEEP Consortium for their support.

