## Supplementary material for "CRUP: A comprehensive framework to predict condition-specific regulatory units"

### Supporting Information (SI)

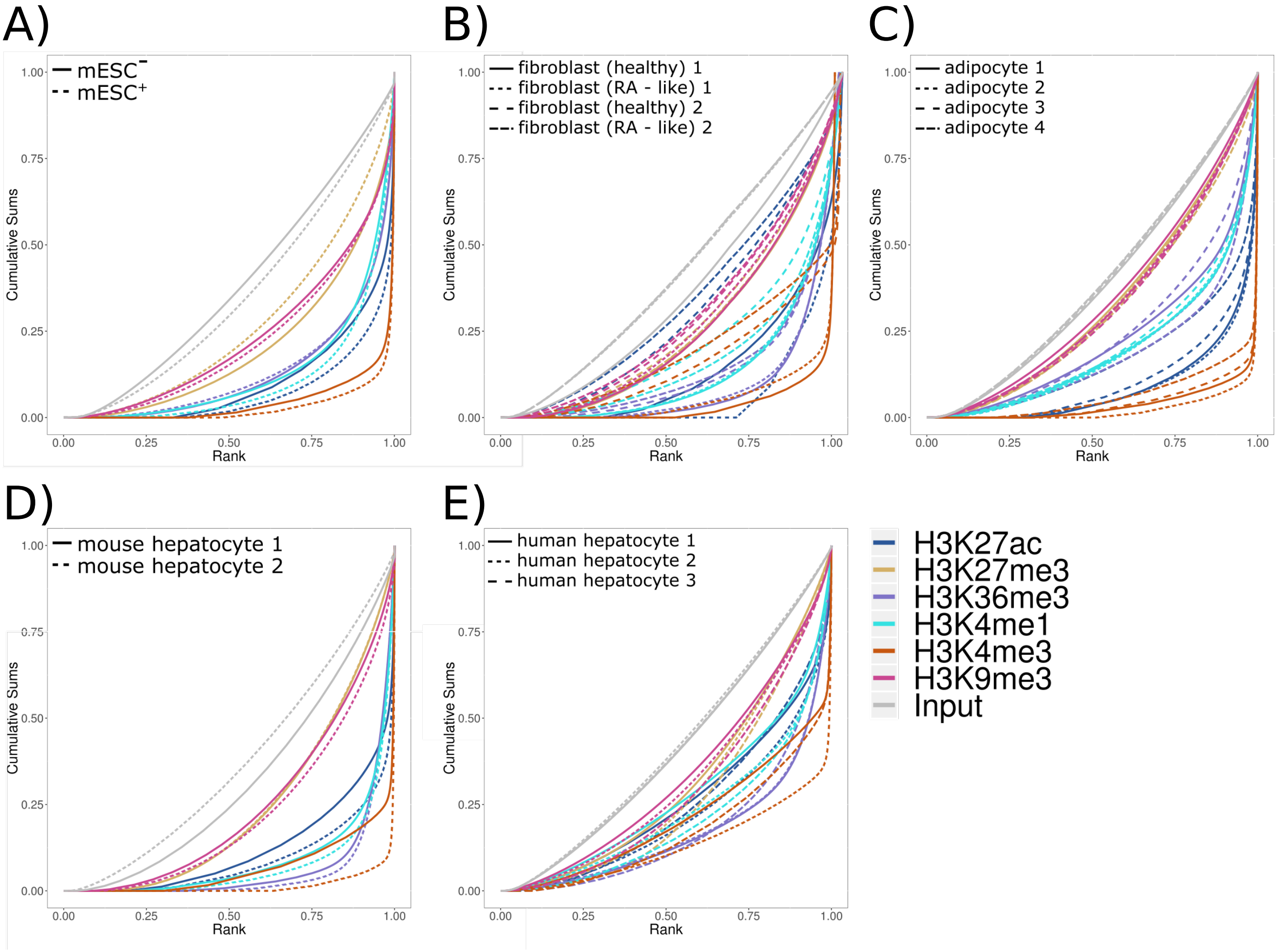


Figure S1: **Fingerprint quality control metrics for ChIP-seq experiments.** Reads with a mapping quality of at least 30 are counted for all adjacent 500bp bins and the cumulative sums are plotted according to their sorted ranks. An ideal input represents a perfect uniform distribution of reads along the whole genome and would generate a straight diagonal line. This is accomplished in all of the analyzed input ChIP-seq experiments (gray lines). For HMs with a very specific and strong ChIP enrichment, a prominent and steep rise of the cumulative sum towards the highest rank would be expected. Most of the profiles show that the analyzed ChIP-seq data have a very good quality. One exception can be observed for the H3K27ac profile generated for the sample 'fibroblast 1' in panel B (blue dashed line). Here it can be seen that nearly 75% have not been sequenced, as almost 75% of all the genomic bins contain 0 reads.

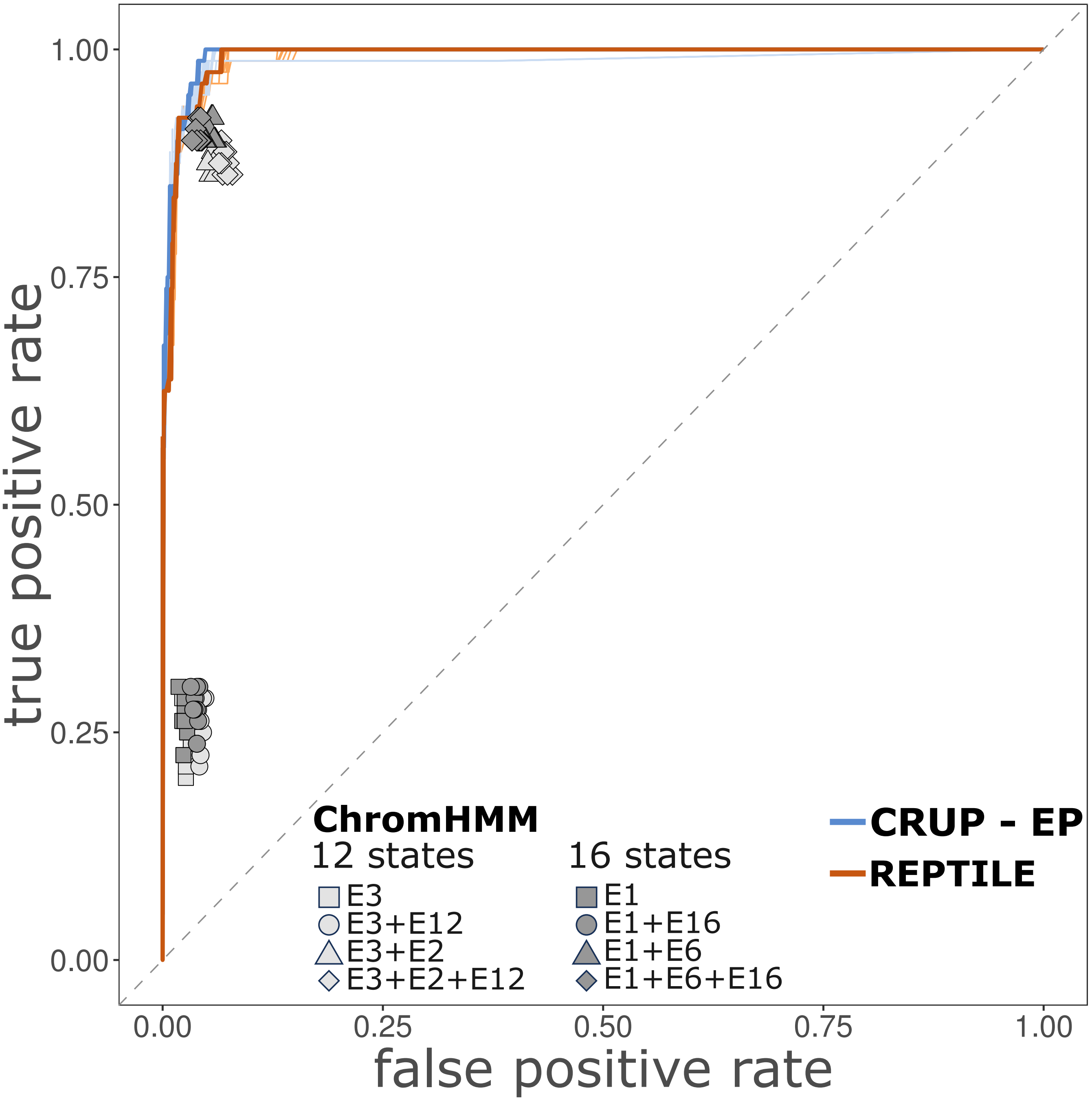


Figure S2: **ROC curve performance of enhancer classifiers in murine ESC.** ROC curves for CRUP-EP (light blue lines) and REPTILE (light orange lines) trained on an mESC sample (mESC$^{+}$) and tested on ten randomly sampled independent test sets. The curves for the best performances are highlighted in darker colors. Additionally, the performance results of different ChromHMM segmentations for the same ten test sets are depicted (gray shapes).

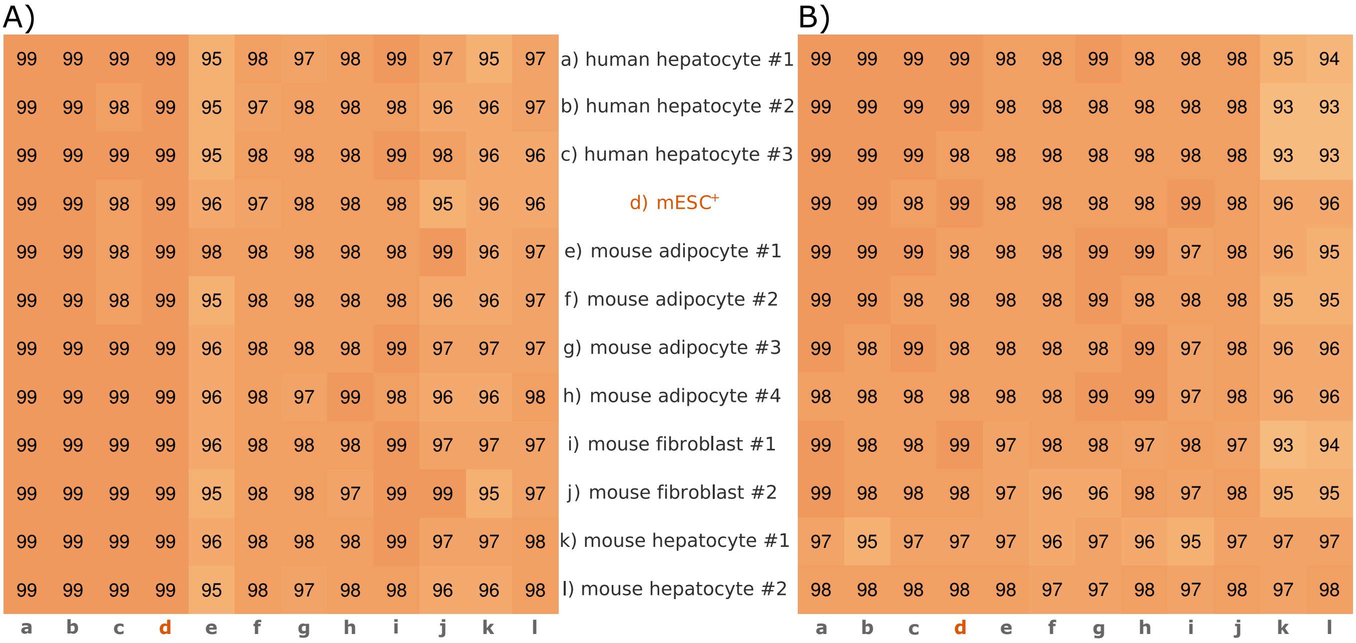


Figure S3: **AUC-ROC performance for predictions across cell lines and species.** Classifiers were trained on and applied to samples from different cell types (hepatocyte, ESC, adipocyte, fibroblast) and species (mouse and human). The results for CRUP-EP (A) and REPTILE (B) can be summarized in 12 $\times$ 12 heatmaps where each entry is shaded according to the computed AUC-ROC (in percent). The origin of the training data a)-l) can be found in the rows and the origin of test sets **a-l** in the columns.

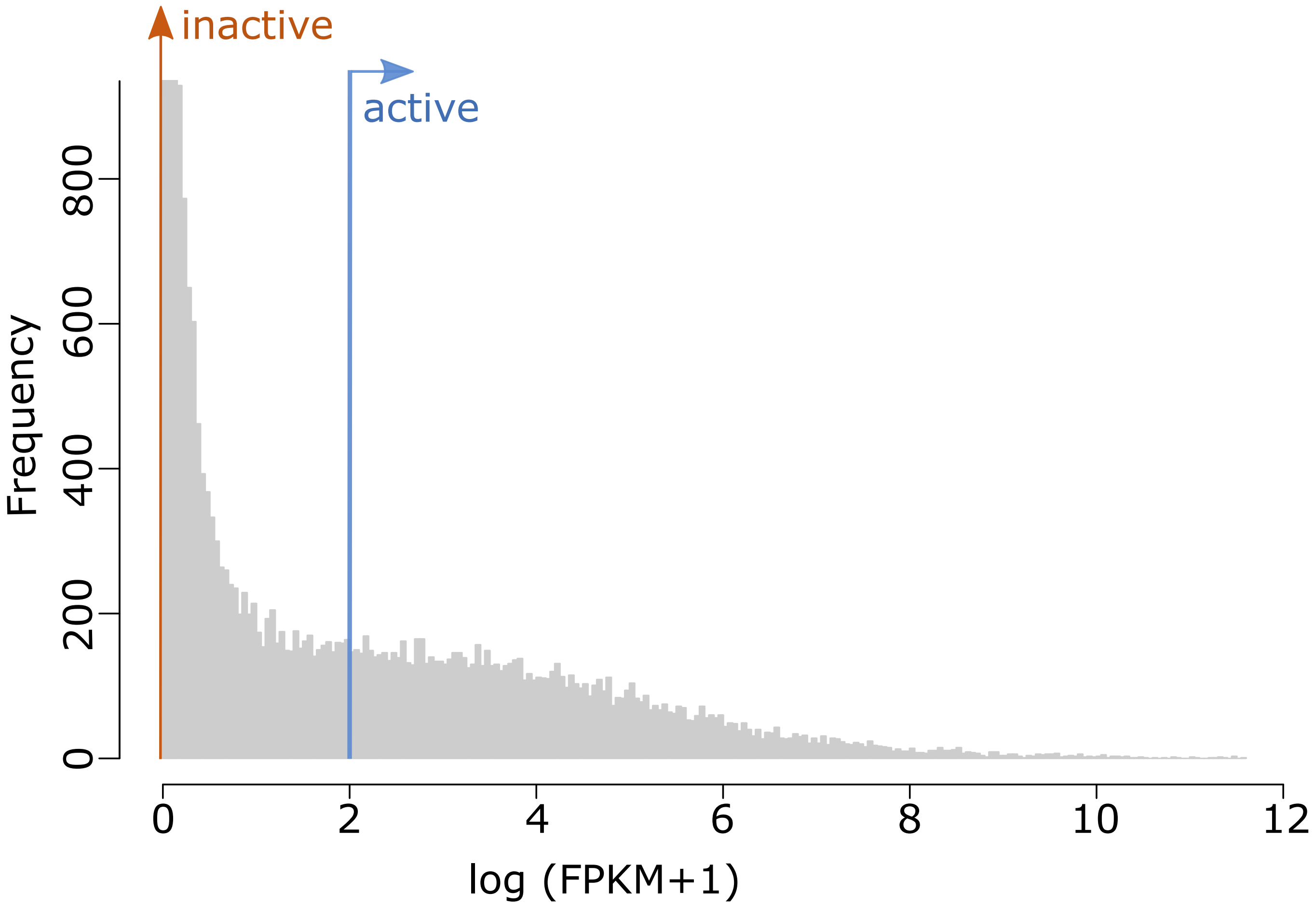


Figure S4: **Distribution of gene expression values in mESC.** Based on RNA-seq data for $mESC^+$ we computed FPKM normalized gene expression values for each gene (gray distribution). We defined a gene with an FPKM value $> 2$ as active (blue), and every gene with FPKM value of zero as inactive (orange).

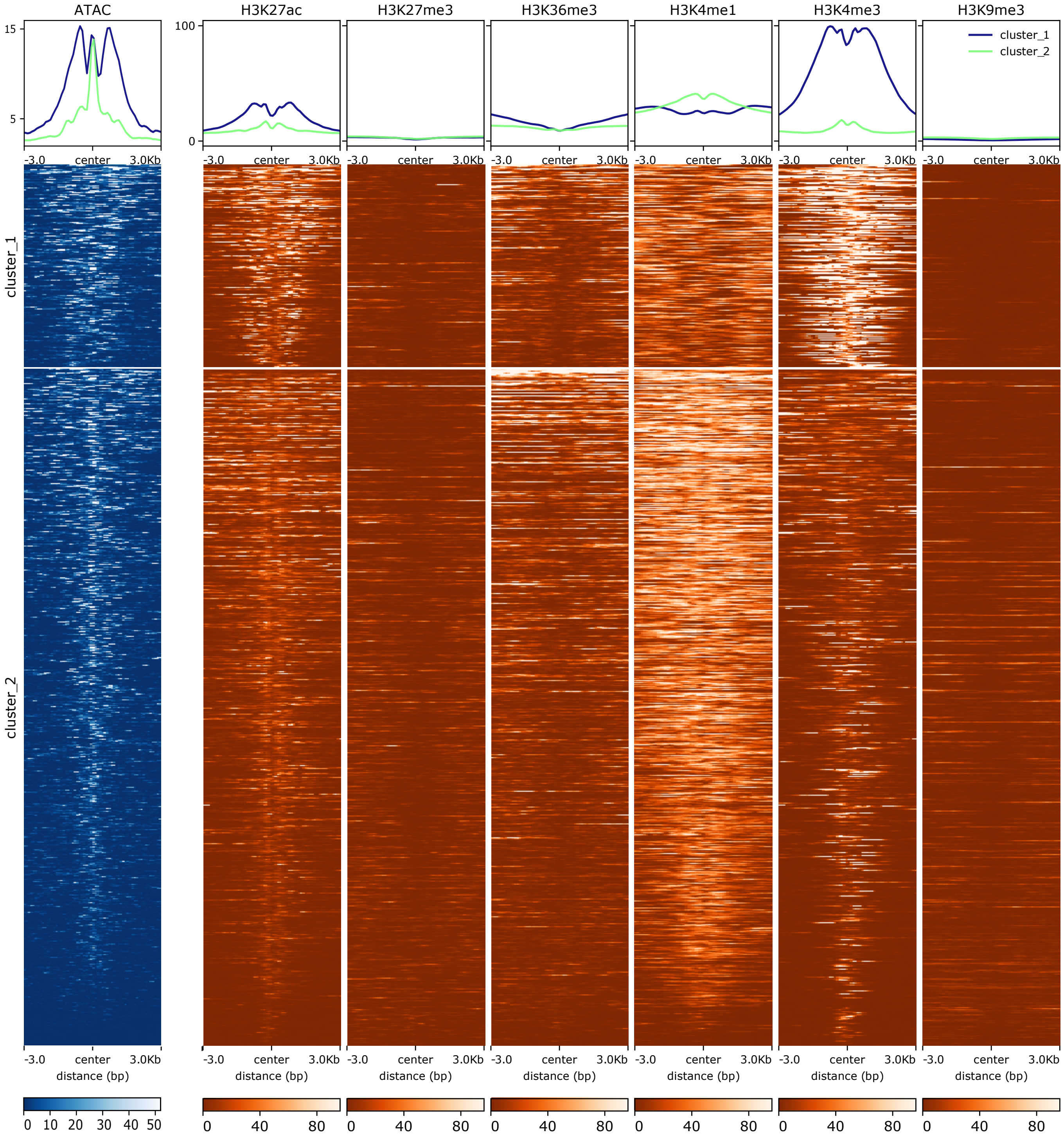


Figure S5: **Count distributions for HMs and open chromatin for predicted enhancers in mESC.** Distributions of HM ChIP-seq and ATAC-seq read counts for 42,530 called enhancer peaks in $mESC^{+}$. A k-means clustering reveals two distinct groups (cluster 1 and 2). Whereas the larger second cluster shows a typical enhancer state with high values for H3K4me1 and H3K27ac, cluster 1 seems to represent promoter-proximal enhancer regions with an additional high enrichment for H3K4me3 and a non-centered ATAC-seq enrichment.

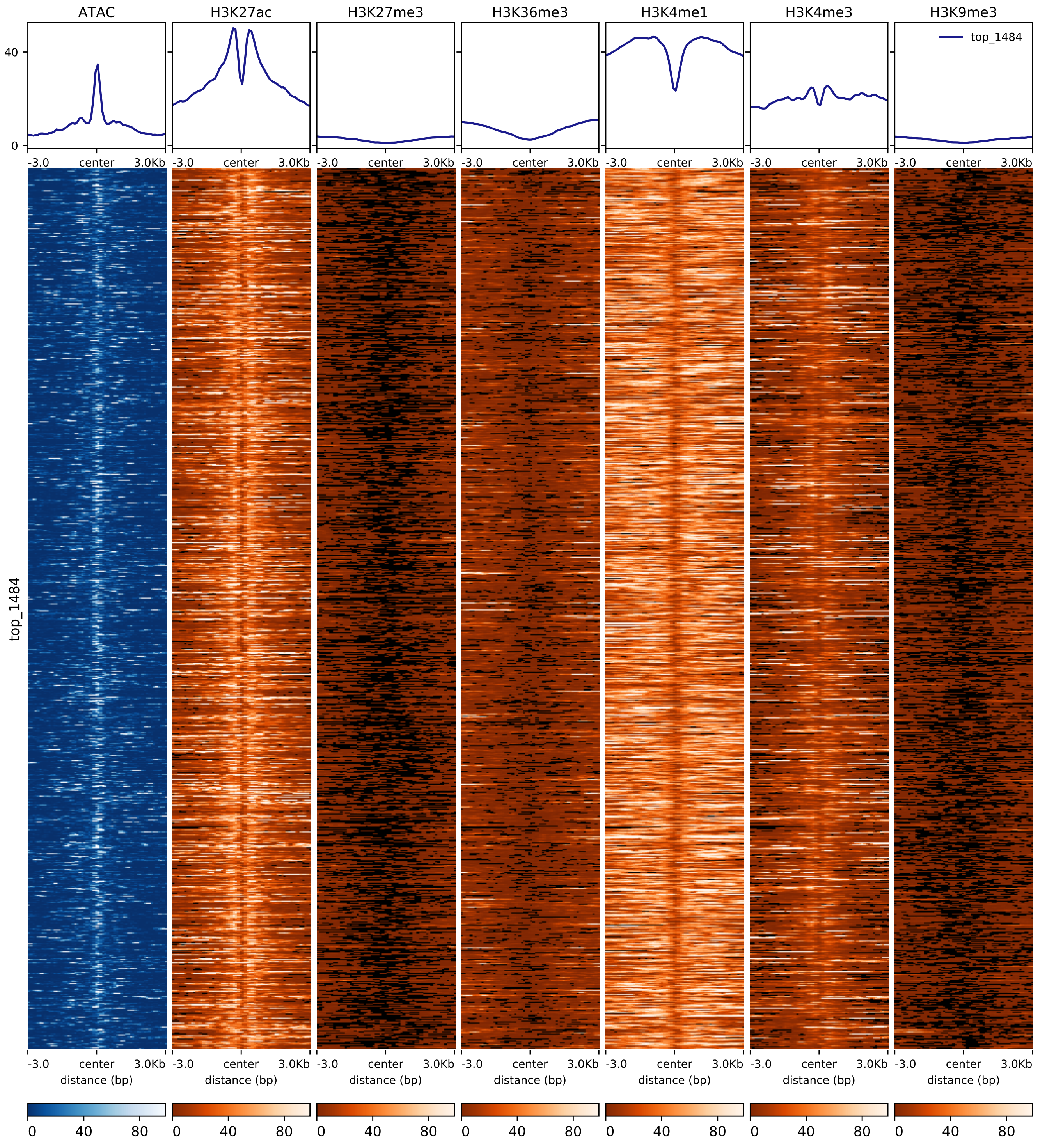


Figure S6: **Count distributions for HMs and open chromatin for top-ranked predicted enhancers in mESC.** Distributions of HM ChIP-seq and ATAC-seq read counts for the top-ranked $1,484$ called enhancer peaks in $mESC^{+}$.

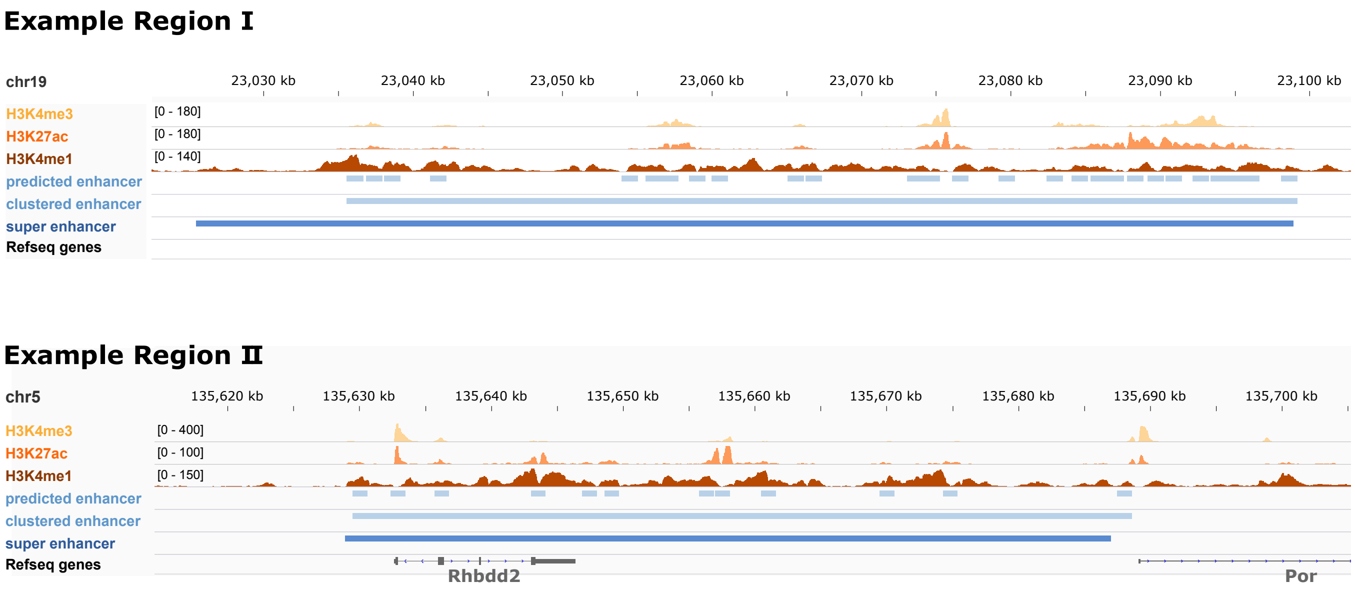


Figure S7: **Two example regions of enhancer clusters in mESC.** Two enhancer clusters in mESC (light blue) coninciding with *'super enhancers'* (dark blue) defined in Novo *et al.* (2018).

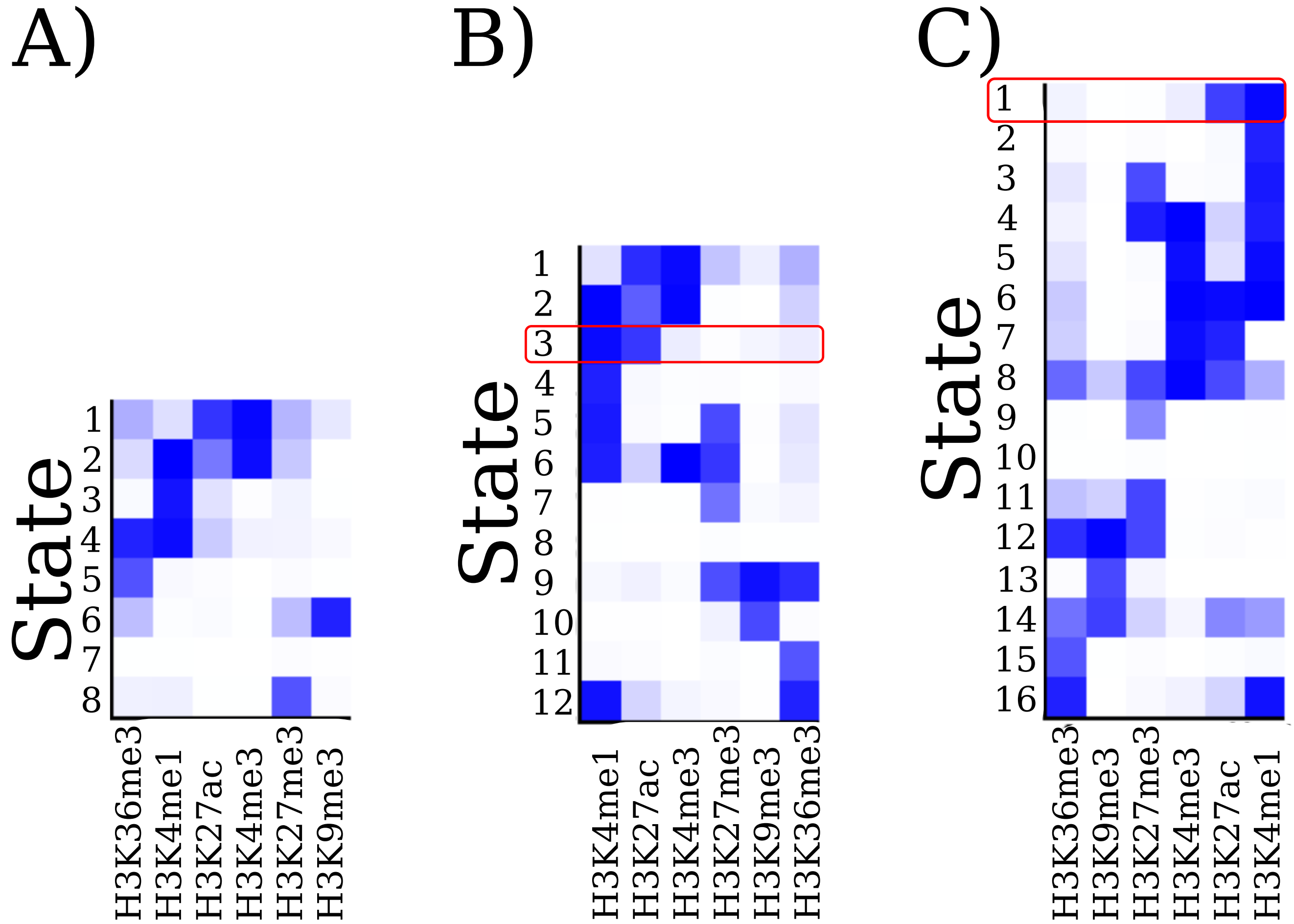


Figure S8: **ChromHMM emission probabilities for mESC data.** ChromHMM was applied on an undifferentiated mESC sample (mESC$^+$) with three different chromatin segmentation states: A) 8 states, B) 12 states and C) 16 states. The heatmaps show the emission probabilities in each defined ChromHMM state, i.e., the probabilities with which each HM is found in each state. According to the combination of H3K27ac and H3K4me1 emission probabilities, we defined the enhancer states for each of the three segmentations. **A)** We could not clearly distinguish an enhancer state from the promoter state (high H3K27ac and high H3K4me3 emission probabilities). **B)** We defined state 3 (highlighted in red) as an enhancer state. The optimal performance results over 10 different test seeds for this ChromHMM setting were: TPR= 0.24, FPR= 0.012, precision=0.66. We also tried to expand this choice by additionally adding state 12 to the enhancer state, which somewhat increased the TPR, but led to worse results regarding the other performance measures: TPR= 0.25, FPR= 0.018, precision= 0.57. **C)** We defined state $E1$ as our enhancer state (highlighted in red) which yielded TPR= 0.29, FPR= 0.012, precision= 0.73 over 10 different test seeds. Also here, including an additional state 16 to the enhancer state definition increased the TPR, but led to worse results regarding the other performances: TPR= 0.32, FPR= 0.02, precision= 0.56.

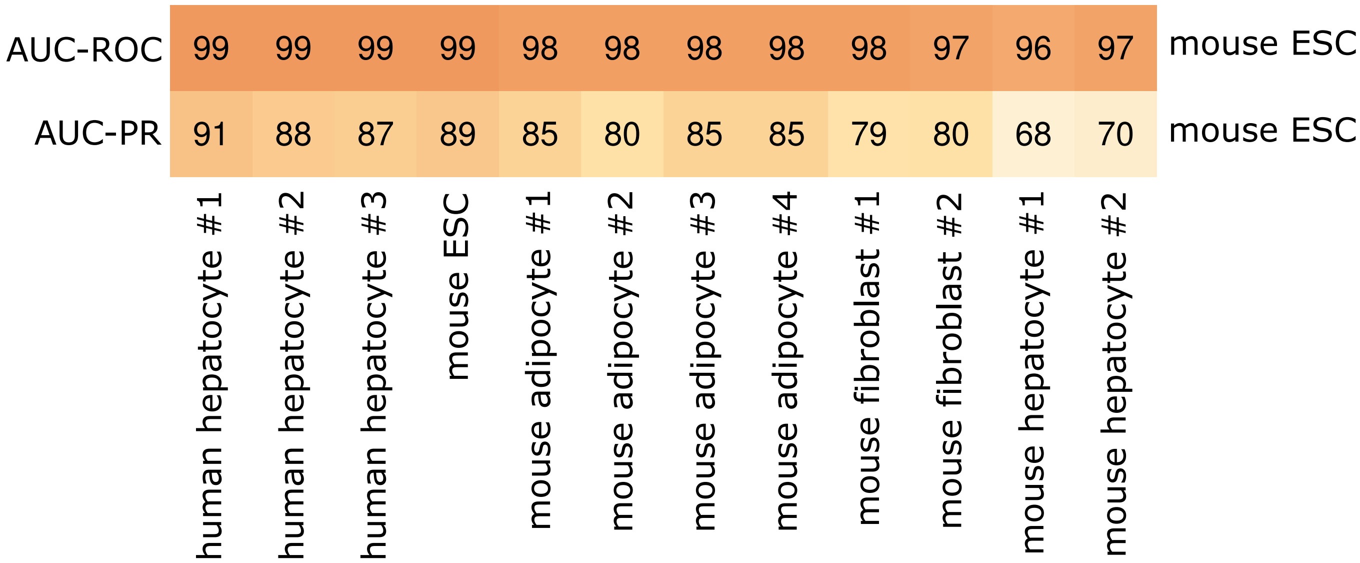


Figure S9: **AUC-ROC and AUC-PR for REPTILE on original training and data set.** We trained a REPTILE classifier on the original p300-based training set using the original HM ChIP-seq data in mESC$^+$ and made predictions on a test set in 12 samples from different cell lines and species. The REPTILE classifier from this setting works worse then using our training and data set.

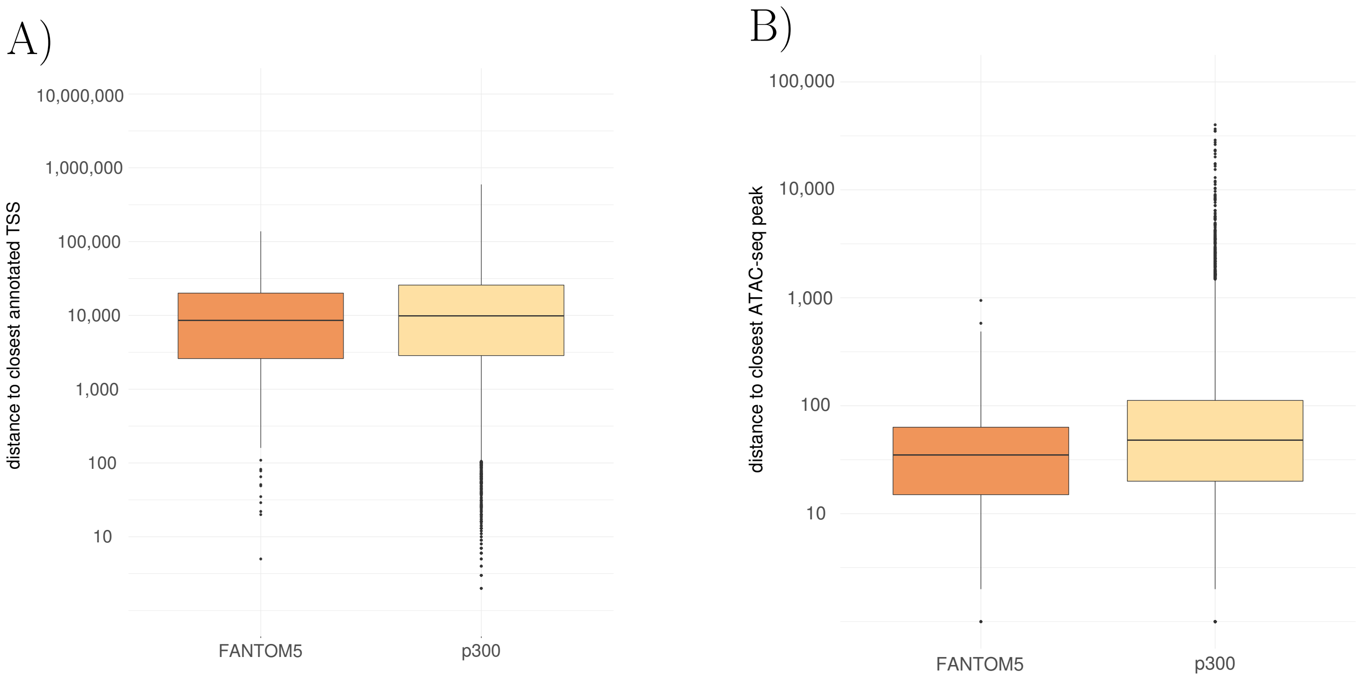


Figure S10: **Distance of FANTOM5 and p300 training sets to closest TSS and ATAC-seq peak in mESC.** We computed the distance of all FANTOM5 enhancers used for training in mESC$^+$ (280 regions) and all p300 ChIP-seq peaks from He *et al.* (2017) (5000 regions) to **A)** the closest annotated transcription start site (TSS) defined by ENSEMBL (GRCm38.90) and **B)** the closest ATAC-seq peak.

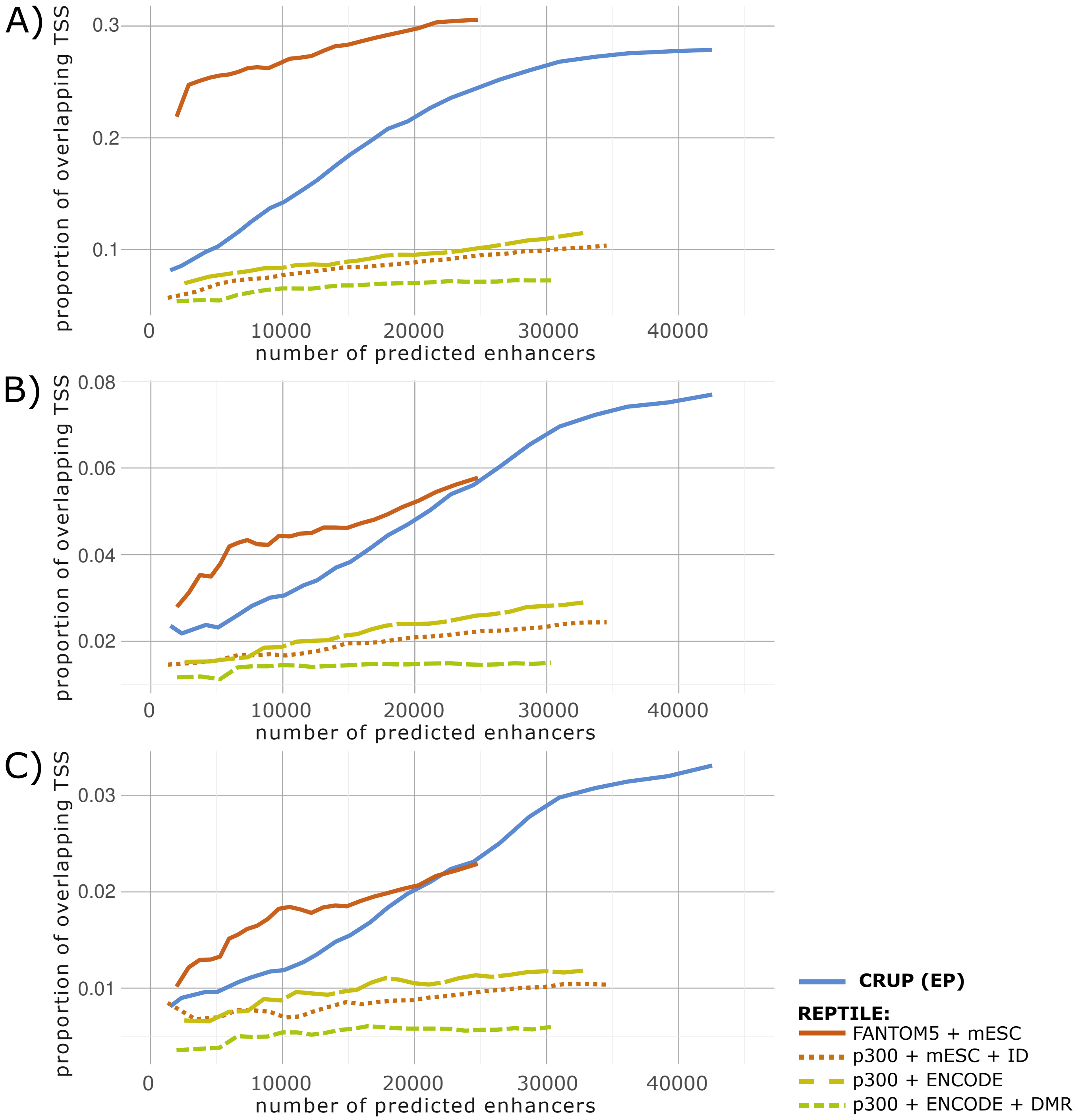


Figure S11: **Proportion of predicted enhancers overlapping TSSs**. We trained our classifier, CRUP-EP, on the optimal setting and the REPTILE classifier for different settings and made genome-wide predictions in mESC$^{+}$. We called enhancers by increasing the probability cutoff and computed the overlap between the summit and annotated transcription start sites (TSSs) defined by ENSEMBL (GRCm38.90).
**A)** TSS plus minus 1000 bp **B)** TSS plus minus 250 bp **C)** TSS plus minus 100 bp

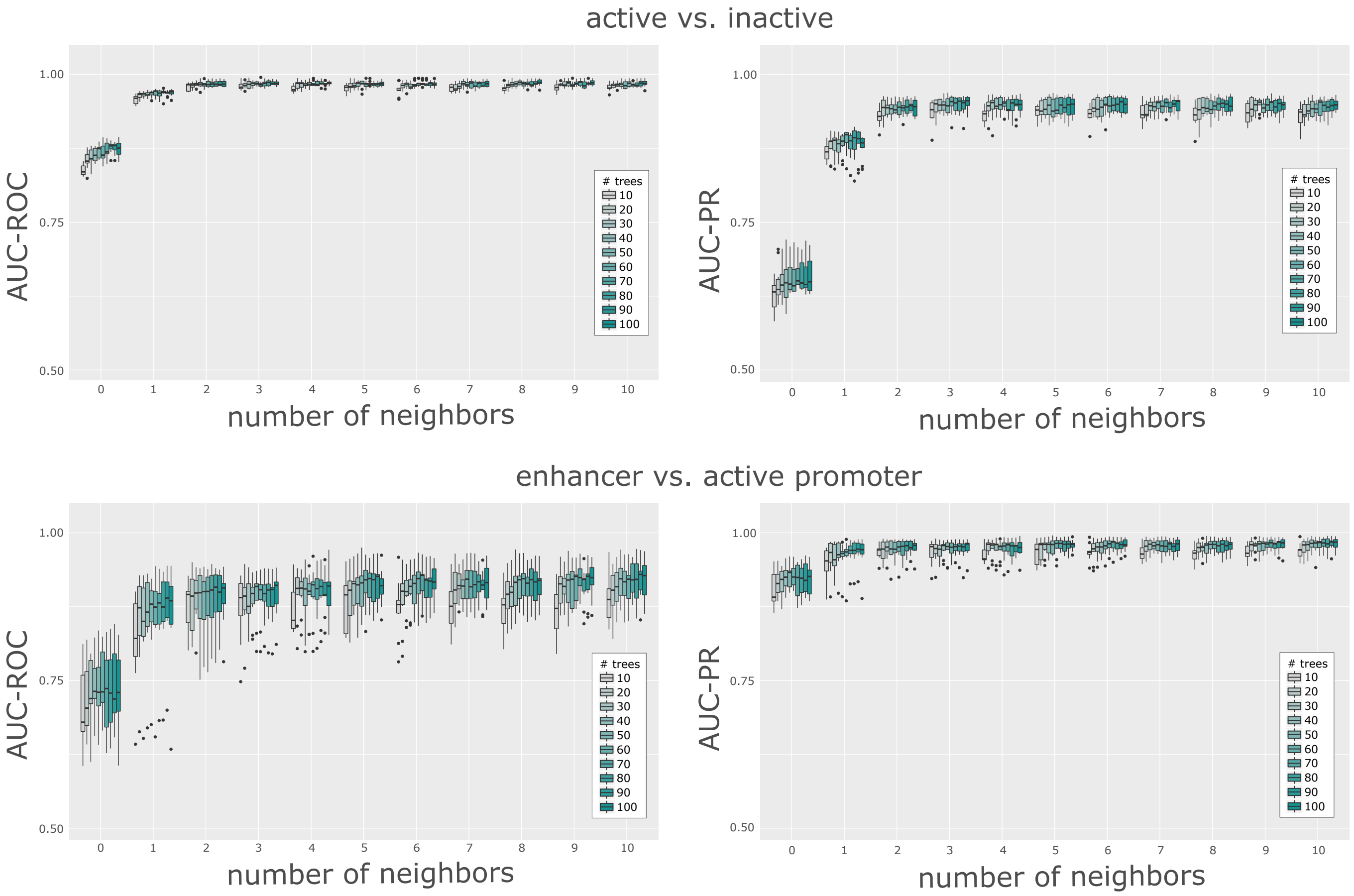


Figure S12: **AUC-ROC and AUC-PR cross-validation results for parameter tuning in mESC.** Using 5-fold cross-validation we evaluated the enhancer classifier embedded in CRUP for 1100 different parameter settings in total which were derived from different choices for training seeds, number of neighboring windows and number of decision trees: $10 \cdot 11 \cdot 10 = 1100$. Based on the results we chose 5 neighboring windows and 70 trees for both classifiers.

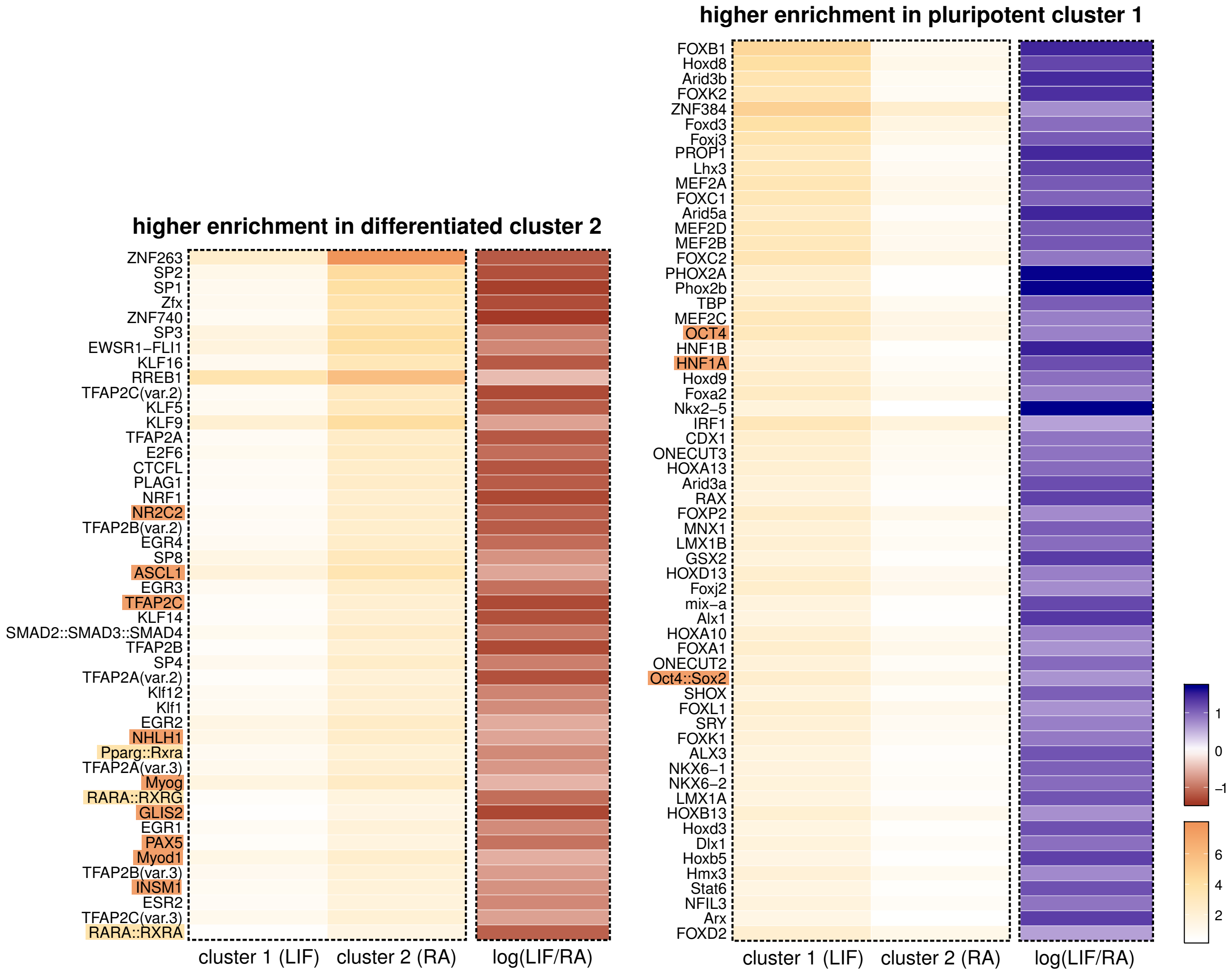


Figure S13: **Motif enrichment for differential enhancers associated with retinoic acid signaling.** We computed the motif enrichment for the differential enhancers which were grouped in cluster 1 (LIF; active enhancers only in the pluripotent state) and cluster 2 (RA; active enhancers only in RA-induced differentiation state). We further filtered for TFs which had an enrichment value of at least 1 in one of the clusters and in addition a difference in enrichment of at least 1 between both clusters, resulting in 106 TFs. Depicted are enrichment values for the two clusters and additionally, the log-fold enrichment between cluster 1 and cluster 2. Red log-fold values indicate TFs with a higher motif enrichment in the RA-cluster (on the left) while TFs with a higher enrichment in the pluripotent cluster 1 result in a blue colour code. TFs enriched in cluster 1 which are marked in orange are part of signaling pathways regulating pluripotency in stem cells. For TFs enriched in cluster 2, the ones highlighted in yellow are either RA receptors or originating form heterodimers with RA receptors. Further functional annotation analysis was used to assign TFs enriched in cluster 2 to *differentiation* and/or *developmental protein* funtional categories which are highlighted in orange (Huang and Lempicki, 2009; Huang *et al.*, 2009).

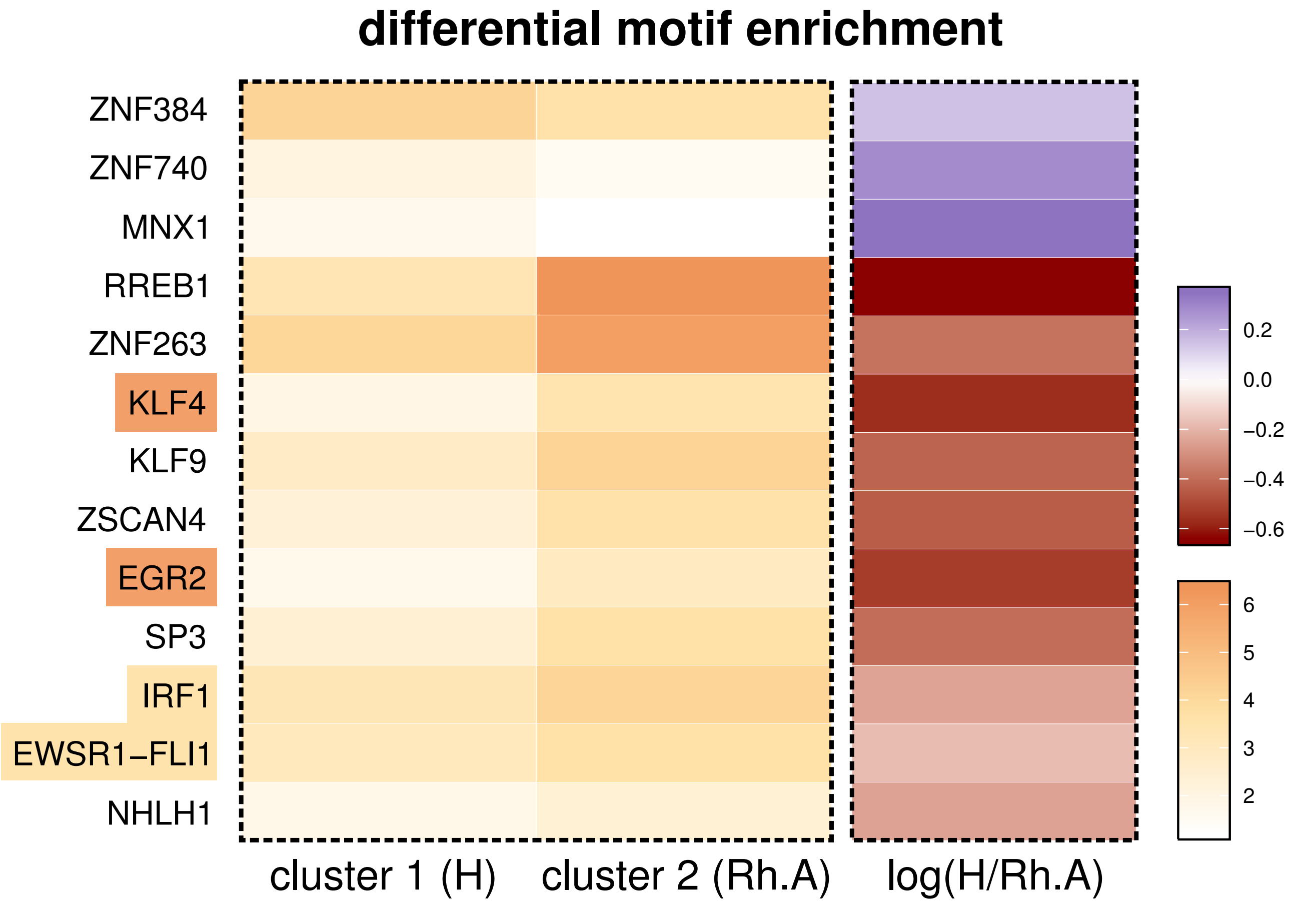


Figure S14: **Motif enrichment for differential enhancers associated with desctructive arthritis in mice.** We computed the motif enrichment for the differential enhancers which were grouped in cluster 1 (H; active enhancers only in healthy samples) and cluster 2 (RH.A; active enhancers only in destructive arthritis samples). We further filtered for TFs which had an enrichment value of at least 1 in one of the clusters and in addition a difference in enrichment of at least 0.5 between both clusters, resulting in 13 TFs. Depicted are enrichment values for these TFs with a specifically high enrichment in cluster 1 or cluster 2, coloured by enrichment. We also computed the log-fold values between the enrichment of the two clusters, where red colours indicate a higher motif enrichment in cluster 2, and blue colours a higher enrichment in cluster 1. TFs highlighted in orange, *KLF4* and *EGR2*, are known key players in rheumatoid arthritis, either as regulator of proinflammatory signaling (Myouzen *et al.*, 2010) or as a risk factor (Luo *et al.*, 2016). The TFs marked in yellow, *IRF1* and *FLI1*, are connected to chronic inflammatory conditions (Salem *et al.*, 2014) or are regulators of TFs associated with autoimmune disease progression in rheumatoid arthritis (Sato *et al.*, 2014).

Table S1: **Summary of experimental data sources used in this work.** Most of the samples are obtained by the German epigenome programme (DEEP) and just two samples were produced in-house (mESC$^{+}$, mESC$^{-}$). Healthy samples obtained by DEEP were used to validate the classification method. Differentiated mouse ESC (mESC$^{-}$) and synovial fibroblast samples, affected by destructive athritis (*'RA - like'*), are further used to identify differentially active enhancers. Additional data was downloaded for as study of mouse neural differentiation (ES, NPC, CN) as well as data for eight different time points in mouse embry midbrain development (Day10.5 to Day0 after birth).

| Abbreviation | Species | Tissue/Cell Type | Project/Study | Condition/Treatment |
|---|---|---|---|---|
| mESC$^{+}$ | mouse | ESC blastocyst | in-house | LIF |
| fibroblast (healthy) | mouse | synovial fibroblast | DEEP | healthy/ no treatment |
| mESC$^{-}$ | mouse | ESC blastocyst | in-house | -LIF, +RA |
| fibroblast (RA - like) | mouse | synovial fibroblast | DEEP | rheumatoid arthritis like |
| adipocyte | mouse | adipocyte/white fat cell | DEEP | healthy/ no treatment |
| mouse hepatocyte | mouse | liver hepatocyte | DEEP | healthy/ no treatment |
| human hepatocyte | human | liver hepatocyte | DEEP | healthy/ no treatment |
| midbrain Day10.5 | mouse | midbrain | Gorkin *et al.* | Day10.5 |
| midbrain Day11.5 | mouse | embryo midbrain | Gorkin *et al.* | Day11.5 |
| midbrain Day12.5 | mouse | embryo midbrain | Gorkin *et al.* | Day12.5 |
| midbrain Day13.5 | mouse | embryo midbrain | Gorkin *et al.* | Day13.5 |
| midbrain Day14.5 | mouse | embryo midbrain | Gorkin *et al.* | Day14.5 |
| midbrain Day15.5 | mouse | embryo midbrain | Gorkin *et al.* | Day15.5 |
| midbrain Day16.5 | mouse | embryo midbrain | Gorkin *et al.* | Day16.5 |
| midbrain Day0 (AB) | mouse | embryo midbrain | Gorkin *et al.* | Day0 after birth |
| ES | mouse | embryonic stem cell | Bonev *et al.* | undifferentiated |
| NPC | mouse | neural progenitor | Bonev *et al.* | neural progenitor |
| CN | mouse | cortical neuron | Bonev *et al.* | cortical neuron |

Table S2: **Overview of sample and replicate size per experimental method.** RNA-seq, ChIP-seq and DNase-seq experiments were used in various steps in our framework. Additionally, ATAC-seq and Hi-C experiments were used for validation. Shown are the number of samples per experiments with the number of replicates in brackets.

| Abbreviation | # RNA-seq | # DNase-seq | # ChIP-seq | # ATAC-seq | # Hi-C |
|---|---|---|---|---|---|
| $mESC^+$ | 1 (3) | 1 (2) | 1 | 1 | - |
| $mESC^-$ | 1 (3) | - | 1 | 1 | - |
| fibroblast (healthy) | 2 | 2 | 2 | - | - |
| fibroblast (RA - like) | 2 | 2 | 2 | - | - |
| adipocyte | - | 1 | 4 | - | - |
| mouse hepatocyte | - | 2 | 2 | - | - |
| human hepatocyte | - | 3 | 3 | - | - |
| midbrain Day10.5 | 1(2) | - | 1(2) | - | - |
| midbrain Day11.5 | 1(2) | - | 1(2) | - | - |
| midbrain Day12.5 | 1(2) | - | 1(2) | - | - |
| midbrain Day13.5 | 1(2) | - | 1(2) | - | - |
| midbrain Day14.5 | 1(2) | - | 1(2) | - | - |
| midbrain Day15.5 | 1(2) | - | 1(2) | - | - |
| midbrain Day16.5 | 1(2) | - | 1(2) | - | - |
| midbrain Day0 (AB) | 1(2) | - | 1(2) | - | - |
| ES | 1(2) | - | 1(2) | - | 1 |
| NPC | 1 | - | 1 | - | 1 |
| CN | 1(2) | - | 1(2) | - | 1 |

Table S3: **Enhancer regions defined by FANTOM5.** CAGE count data was downloaded for mouse embryonic stem cells, mouse synovial fibroblasts,mouse adipocytes and mouse and human hepatocytes. Depending on the available number of replicates ('$\sum$ *Repl.*') all regions ('# *Regions*') where narrowed down to a set of high confidence enhancers ('*Criterium*'). For example, we used count data of three biological replicates from murine hepatocytes and chose 753 enhancers which had eight and more counts in all three replicates.

| Abbr. | FANTOM5 Cell Line Description | Criterium ($\sum$ Repl.) | # Regions |
|---|---|---|---|
| mESC$^+$ | • ES-OS25 embryonic stem cells, DMSO control<br>• ES-OS25 embryonic stem cells, untreated control<br>• ES-Ert2 embryonic stem cells, untreated control, 48hr<br>• ES-OS25 embryonic stem cells, untreated siRNA control<br>• ES-OS25 embryonic stem cells, scrambled siRNA control | $\geq$ 4 counts in all (13) | 372 |
| adipocyte | • ST2 (mesenchymal stem cells) cells, differentiation to adipocytes, day06 | > 3 counts in all (3) | 756 |
| fibroblast (healthy) | • mouse fibroblast cell line: CRL-1658 NIH/3T3 | $\geq$ 2 counts in any (1) | 683 |
| mouse hepatocyte | • liver sinusoidal endothelial cells, partial hepatectomy, 01week | $\geq$ 8 counts in all (3) | 753 |
| human hepatocyte | • liver, adult, pool1 | $\geq$ 8 counts in any (1) | 298 |

Table S4: **Final enhancer regions defined by FANTOM5 and DNaseI peaks.** DNase-seq peaks were called for each sample/replicate. The overlap of DNAseI peaks and the filtered FANTOM5 regions from Table S1 build the final enhancer lists used in our workflow. Here, for each type of tissue, we chose the overlap set with the maximal size (bold). For example, we used the 239 DNaseI peaks of sample 1 that overlap with FANTOM5 as representative enhancer set for mouse hepatocytes.

| Abbreviation | sample/replicate | # DNaseI peaks | # FANTOM5 | # overlap |
|---|---|---|---|---|
| mESC$^+$ | replicate 1 | 123576 | 372 | **280** |
| | replicate 2 | 88973 | | 250 |
| adipocyte | sample 1 | 43814 | 756 | **292** |
| fibroblast (healthy) | sample 1 | 90858 | 683 | **251** |
| | sample 2 | 65682 | | 141 |
| mouse hepatocyte | sample 1 | 51110 | 753 | **239** |
| | sample 2 | 44336 | | 227 |
| human hepatocyte | sample 1 | 86296 | 298 | **217** |
| | sample 2 | 44290 | | 176 |
| | sample 3 | 40438 | | 176 |

Table S5: **Active promoter regions defined by RNA-seq cutoff and DNase peaks.** We expanded the TSSs of active genes (”active” according to definition in Section 5.8) symmetrically to a total length of 100 bp and computed the overlap with DNAse summits in the same tissue (not always the same sample). Only expanded TSSs containing a DNase summit are finally used to define active promoters.

| sample/replicate | # active promoter | DNaseI sample | # overlap |
|---|---|---|---|
| mESC$^+$ | 10,044 | replicate 1 | 2,853 |
| adipocyte sample 1 | 9,217 | sample 1 | 2,273 |
| adipocyte sample 2 | 9,206 | sample 1 | 2,295 |
| adipocyte sample 3 | 9,245 | sample 1 | 2,317 |
| adipocyte sample 4 | 9,317 | sample 1 | 2,339 |
| fibroblast (healthy) sample 1 | 10,593 | sample 1 | 2650 |
| fibroblast (healthy) sample 2 | 10,326 | sample 1 | 2528 |
| mouse hepatocyte sample 1 | 8,392 | sample 1 | 2,299 |
| mouse hepatocyte sample 2 | 8,417 | sample 1 | 2,318 |
| human hepatocyte sample 1 | 6,668 | sample 1 | 1,689 |
| human hepatocyte sample 2 | 6,853 | sample 1 | 1,749 |
| human hepatocyte sample 3 | 7,021 | sample 1 | 1,670 |

Table S6: **Regulatory units with enhancers only active in destructive arthritis samples and the top 5 associated KEGG pathways enriched in the putative target genes.**

| KEGG Pathway | differential enhancer | gene ID | gene symbol |
|---|---|---|---|
| Chemokine signaling pathway path:mmu04062 | chr1:128722101-128723300 | 12767 | Cxcr4 |
| | chr3:105894101–105894700 | 109905 | Rap1a |
| | chr4:3680001–3680500 | 17096 | Lyn |
| | chr5:134231901–134233300 | 17969 | Ncf1 |
| | chr9:99182301–99183300 | 74769 | Pik3cb |
| | chr9:123930801–123931300 | 12768 | Ccr1 |
| | chr9:123930801–123931300 | 12770 | Ccr1l1 |
| | chr9:123930801–123931300 | 12771 | Ccr3 |
| | chr9:123930801–123931300 | 12772 | Ccr2 |
| | chr9:123930801–123931300 | 12774 | Ccr5 |
| | chr11:70468401–70469100 | 216869 | Arrb2 |
| | chr11:70468401–70469100 | 66102 | Cxcl16 |
| | chr17:57347001–57348000 | 22324 | Vav1 |
| Fc gamma R-mediated phagocytosis path:mmu04666 | chr1:74374901–74375700 | 76709 | Arpc2 |
| | chr1:87639601–87640400 | 16331 | Inpp5d |
| | chr4:3680001–3680500 | 17096 | Lyn |
| | chr4:129496301–129497000 | 17357 | Marcksl1 |
| | chr5:134231901–134233300 | 17969 | Ncf1 |
| | chr9:99182301–99183300 | 74769 | Pik3cb |
| | chr10:81297201–81299100 | 18717 | Pip5k1c |
| | chr11:70468401–70469100 | 18806 | Pld2 |
| | chr17:57347001–57348000 | 22324 | Vav1 |
| C-type lectin receptor signaling pathway path:mmu04625 | chr1:58726401–58728600 | 12370 | Casp8 |
| | chr1:131084101–131084800 | 16153 | Il10 |
| | chr1:131084101–131084800 | 17164 | Mapkapk2 |
| | chr6:123261101–123261900 | 17474 | Clec4d |
| | chr6:123261101–123261900 | 56619 | Clec4e |
| | chr6:123261101–123261900 | 56620 | Clec4n |
| | chr6:123261101–123261900 | 69810 | Clec4b1 |
| | chr6:129350701–129351500 | 56760 | Clec1b |
| | chr9:99182301–99183300 | 74769 | Pik3cb |
| | chr11:59541201–59543900 | 216799 | Nlrp3 |
| Natural killer cell mediated cytotoxicity path:mmu04650 | chr1:171843101–171844100 | 12506 | Cd48 |
| | chr1:171843101–171844100 | 18106 | Cd244 |
| | chr7:30393201–30394600 | 22177 | Tyrobp |
| | chr7:30393201–30394600 | 23900 | Hcst |
| | chr7:30419401–30423900 | 22177 | Tyrobp |
| | chr7:30419401–30423900 | 23900 | Hcst |
| | chr9:99182301–99183300 | 74769 | Pik3cb |
| | chr17:57347001–57348000 | 22324 | Vav1 |
| | chr19:34473801–34474700 | 14102 | Fas |
| Kaposi sarcoma-associated herpesvirus infection path:mmu05167 | chr1:58726401–58728600 | 12370 | Casp8 |
| | chr1 131084101–131084800 | 17164 | Mapkapk2 |
| | chr4:3680001–3680500 | 17096 | Lyn |
| | chr5:3500901–3502800 | 12571 | Cdk6 |
| | chr9:99182301–99183300 | 74769 | Pik3cb |
| | chr9:123930801–123931300 | 12768 | Ccr1 |
| | chr9:123930801–123931300 | 12770 | Ccr1l1 |
| | chr9:123930801–123931300 | 12771 | Ccr3 |
| | chr9:123930801–123931300 | 12774 | Ccr5 |
| | chr19:34473801–34474700 | 14102 | Fas |
